# MTARC1 Regulates Lipid Droplet Degradation via Phospholipid Remodeling in Metabolic Fatty Liver Disease

**DOI:** 10.1101/2025.07.17.665462

**Authors:** Meng Tie, Liwei Hu, Yunzhi Yang, Shaoxuan Song, Qihan Zhu, Jun Li, Wenjing Wang, Peng Xu, Juan Yu, Mengyue Wu, Tianheng Zhao, Delong Yuan, Hongyu Bao, Xiuyun Wang, Irfan J. Lodhi, Yong Chen, Yali Chen, Anyuan He

## Abstract

Metabolic dysfunction-associated steatotic liver disease (MASLD) is a spectrum of liver disease, including simple fatty liver, hepatic steatosis, fibrosis, cirrhosis, and hepatocellular carcinoma, with limited treatment options. *MTARC1* p.A165T variant is associated with reduced risk of MASLD. We previously showed that this variant promotes MTARC1 protein degradation, suggesting MTARC1 inactivation may represent a potential therapeutic strategy. Here, we report that global or liver specific *Mtarc1* knockout markedly suppresses diet-induced hepatic TG accumulation, liver injury, inflammation, and fibrosis in a manner dependently on lipolysis and lipophagy. MTARC1 inactivation post-transcriptionally upregulates PEMT and CEPT1 expression, consequently elevating GPL levels, including PC, PE, and lyso-PE, three major GPLs of hepatic LDs. Furthermore, we observed that MTARC1-deficient hepatocytes exhibited smaller but more numerous LDs compared to controls, under comparable cellular TG levels during lipolysis/lipophagy inhibition. Importantly, inhibition of CEPT1 or PEMT could attenuated the hepatoprotective effect of MTARC1 deficiency. Mechanistically, subcellular GPL remodeling induced by MTARC1 deficiency reduce LD size while increase their surface-to-volume ratio, which in turn promote TG degradation through lipolysis and lipophagy. Collectively, our findings identify MTARC1–GPL biosynthesis–LD degradation axis as a key regulator of fatty liver disease and highlight MTARC1 inhibition as a promising therapeutic strategy for MASLD.

## Introduction

Metabolic dysfunction-associated steatotic liver disease (MASLD), previously termed as nonalcoholic fatty liver disease (NAFLD), is a spectrum of liver disease, including simple fatty liver, hepatic steatosis, fibrosis, cirrhosis, and hepatocellular carcinoma ^1,2^. MASLD incidence continues to rise and affects nearly 1 billion people globally, with limited treatment options. The mitochondrial amidoxime-reducing component (MTARC1) represents as a potential therapeutic target for treating fatty liver disease, as human genetic studies have revealed that a set of *MTARC1* variants is associated with all-cause fatty liver disease ^3^. *MTARC1* p.A165T, a hepatoprotective variant with 29% minor allele frequency, promotes the degradation of MTARC1 protein, documented by us and others ^4–8^. MTARC1 is a Mo (molybdenum)-containing enzyme known to activate N-hydroxylated prodrugs ^9–11^. Although efforts have been made to determine the role of MTARC1 in fatty liver disease ^12, 13^, the mechanism through which MTARC1 affects lipid metabolism and thereby modulates MASLD progression remains to be elucidated.

The excessive accumulation of triglycerides (TGs), the predominant neutral lipids in hepatocytes, is a pathological hallmark of fatty liver disease, thus rendering TG-lowering approaches viable therapeutic strategies. Intracellular TG stored in LDs undergoes mobilization through two principal pathways: lipolysis and lipophagy. The lipolytic cascade involves sequential hydrolysis by adipose triglyceride lipase (ATGL), hormone-sensitive lipase (HSL), and monoglyceride lipase (MGL). Lipophagy, the selective autophagic degradation of LDs, relies on lysosomal acid lipase (LAL)-mediated TG hydrolysis to release fatty acids. Substantial evidence demonstrates that both direct and indirect activation of these pathways could ameliorate fatty liver disease^14–17^.

Phosphatidylcholine (PC) and phosphatidylethanolamine (PE) are the two major classes of glycerophospholipids (GPLs). PC can be synthesized from choline via the CDP-choline pathway, which is mediated by two enzymes: choline/ethanolamine phosphotransferase 1 (CEPT1)—an ER-localized enzyme, and choline phosphotransferase 1 (CPT1)—a Golgi-localized enzyme ^18^. In the liver, PC can also be synthesized from PE via the phosphatidylethanolamine N-methyltransferase (PEMT)-mediated methylation pathway. PE is synthesized via two major pathways in mammalian cells: the CDP-ethanolamine pathway and the phosphatidylserine decarboxylase (PSD) pathway ^18^. Notably, these two PE synthesis pathways occur in spatially distinct organelles—the ER (CDP-ethanolamine pathway) and mitochondria (PSD pathway) ^18^. Importantly, GPL homeostasis is tightly regulated at both cellular and subcellular levels. In hepatocytes, GPLs play critical roles in key lipid metabolism processes, including very-low-density lipoprotein (VLDL) assembly and lipid droplet (LD) dynamics.

Here, we report that global or liver specific *Mtarc1* knockout markedly suppresses diet-induced hepatic TG accumulation, liver injury, inflammation, and fibrosis. MTARC1 deficiency’s hepatoprotection require both lipolysis and lipophagy. Interestingly, our study indicates that MTARC1 inactivation post-transcriptionally upregulates PEMT and CEPT1 expression without affecting their protein stability, consequently elevating GPL levels, including PC, PE, and LPE, three major GPLs of hepatic LDs. Furthermore, we observed that MTARC1-deficient hepatocytes exhibited smaller but more numerous LDs compared to controls, under comparable cellular TG levels during lipolysis/lipophagy inhibition. Importantly, inhibition of CEPT1 or PEMT could attenuated the hepatoprotective effect of MTARC1 deficiency, establishing a causal link between GPL remodeling and the observed phenotype. Mechanistically, subcellular GPL remodeling induced by MTARC1 deficiency reduce LD size while increase their surface-to-volume ratio, which in turn promote TG degradation through lipolysis and lipophagy. Collectively, our findings identify MTARC1–GPL biosynthesis–LD degradation axis as a key regulator of fatty liver disease and highlight MTARC1 inhibition as a promising therapeutic strategy for MASLD.

## Results

### Mtarc1 global knockout mice are protected from diet-induced fatty liver disease

*MTARC1* p.A165T variant is associated with a lower risk of MAFLD. We and others have proved that this variant promotes the degradation of MTARC1 protein ^4–6^, leading us to hypothesize that MTARC1 inactivation might represent a viable therapeutic strategy for fatty liver disease. To address this hypothesis, a mouse model with global deletion of *Mtarc1* was generated (Figure S1A). Genotyping by PCR and sanger sequencing confirmed successful knockout (Figure S1B to D). The mutant mice were born at the expected mendelian frequency and displayed no overt abnormalities.

Hepatic qPCR assay verified the absence of *Mtarc1* messenger from homozygotes without any effect on the expression of *Mtarc2* encoding MTARC2, another Mo-containing enzyme (Figure S1E). Notably, while heterozygous mice showed significantly reduced hepatic *Mtarc1* mRNA levels compared to wild-type (Figure S1E), their MTARC1 protein levels remained unaffected (Figure S1F). Therefore, we used heterozygotes (control) and homozygotes (MKO) for subsequent experiments for the convenience of animal breeding.

We next subjected male control and MKO mice to a choline-deficient, L-amino acid-defined high-fat diet (CDAHFD), an established model diet for inducing fatty liver disease (Figure 1A). After 8 weeks of feeding, while body weights remained comparable between genotypes (Figure S2A), MKO mice exhibited significantly lower liver-to-body weight ratios (Figure 1B). Biochemical analysis revealed a selective reduction in hepatic triglycerides (TGs) but not cholesterol in MKO mice (Figure 1C and Figure S2B), which was corroborated by decreased lipid accumulation in Oil Red O and H&E staining (Figure 1D). The MKO group showed marked improvements in liver injury markers, with lower serum levels of ALT, AST, and total bilirubin (Figure 1E to G). Notably, serum parameters including TGs, total cholesterol, glucose, and NEFA remained unchanged (Figure S2C to F). In line with the reduced liver injury, MKO mice had reduced hepatic expression of *Tgfb1* and *Il1b*, two markers of inflammation, and decreased expression of *Col1a1*, *Spp1*, and *Timp1*, the markers of hepatic fibrosis (Figure 1H). Lower COL1A1 protein levels and reduced Sirius Red staining indicated diminished hepatic fibrosis in MKO mice (Figure 1I, J).

**Fig 1.**
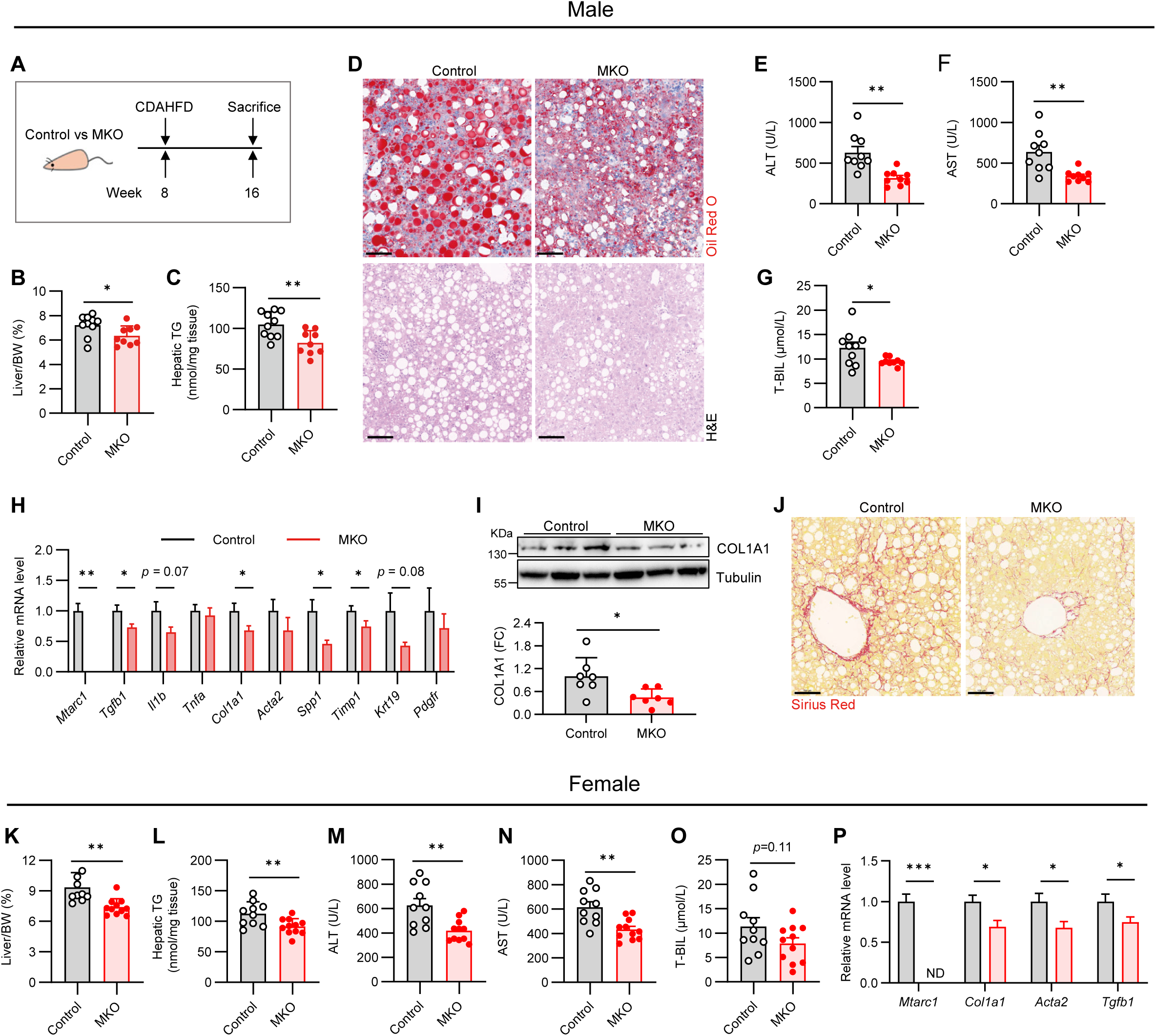
*Mtarc1* global knockout mice are protected from diet-induced hepatic steatosis. (A-J) Male control and *Mtarc1* null mice (MKO) were challenged with CDAHFD for eight weeks and sacrificed for phenotyping (n = 9-10/group). (A) Animal diet challenge experiment workflow. (B) Liver body weight ratio (%). (C) Hepatic TG content. (D) Representative images of livers subjected to Oil Red O staining and H&E staining (Scale bar, 100 μm). (E) Activity of serum alanine aminotransferase (ALT). (F) Activity of serum aspartate aminotransferase (AST). (G) Serum total-bilirubin level. (H) Hepatic qPCR assay for genes as indicated. (I) Immunoblotting for COL1A1 and quantification. (J) Representative images of liver sections subjected to Sirius Red staining (Scale bar, 100 μm). (K-P) Female control and MKO mice were challenged with CDAHFD for eight weeks and sacrificed for phenotyping (n = 10/group). (K) Liver body weight ratio (%). (L) Hepatic TG content. (M) Activity of serum ALT. (N) Activity of serum AST. (O) Serum total-bilirubin level. (P) Hepatic qPCR assay for genes as indicated. Data were expressed as mean ± SEM and analyzed by Student’s t-test. * *p* < 0.05, ** *p* < 0.01.

We subsequently extended the diet challenge experiment in female mice and observed comparable hepatoprotective effects of MTARC1 deficiency (Figure 1K to P, Figure S2G). Body weights, serum TG, cholesterol, and glucose were remained unchanged between two genotypes (Figure S2H to K). Intriguingly, we detected no significant phenotype when a high-fructose/high-fat diet (HFHFD) was employed (Figure S3A to I). Collectively, our data demonstrate that *Mtarc1* knockout confers robust protection against CDAHFD-induced hepatic steatosis, inflammation and fibrosis.

### Mtarc1 liver-specific knockout mice are protected from diet-induced fatty liver disease

Given the established importance of intercellular and interorgan crosstalk in fatty liver disease progression ^19, 20^, we investigated whether inhibiting MTARC1 in a hepatocyte-specific manner could be sufficient to improve hepatic steatosis. To this end, we generated liver-specific *Mtarc1* knockout mice by injecting conditional Cas9 mice with AAV8-TBG-Cre (Figure 2A). This viral vector drives Cas9 expression specifically in hepatocytes, along with two guide RNAs targeting the *Mtarc1* gene. Hepatic *Mtarc1* deletion was verified by qPCR (Figure 2B) and immunoblotting assays (Figure 2C). Of note, MTARC1 share higher similarity with its paralog MTARC2 ^7^. The expression of *Mtarc2* was unaffected by *Mtarc1* deletion (Figure 2B), verifying the fidelity of guide RNAs employed.

**Fig 2.**
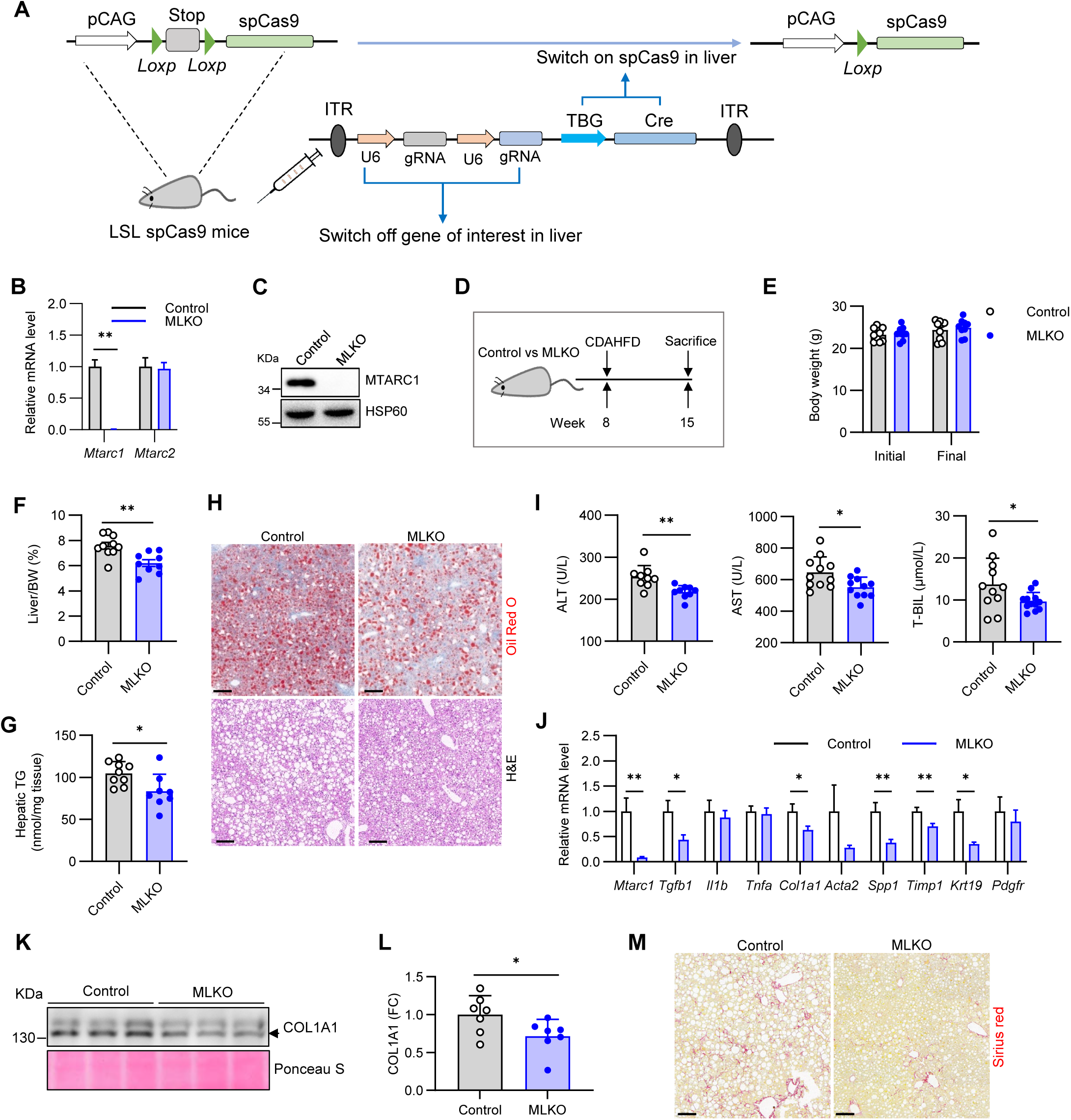
*Mtarc1* liver-specific knockout mice are protected from diet-induced hepatic steatosis. (A) Graphical representation of gene targeting strategy for generation of mice with hepatic deletion of the gene of interest using AAV8 and LSL-Cas9 mice. Briefly, LSL-Cas9 adult mice were tail vein injected with AAV8 carrying either U6.sg1.U6.sg2.TBG.Cre or non-specific control sgRNA.TBG.Cre. (B) qPCR assay for livers from control and MLKO mice (mice with hepatic deletion of *Mtarc1*) (n = 3/group). (C) Immunoblotting for livers from control and MLKO mice. (D-M) Control and MLKO mice were challenged with CDAHFD for seven weeks and sacrificed for phenotyping (n = 8 to 11/group). (D) Animal diet challenge experiment workflow. (E) Initial and final body weights. (F) Liver body weight ratio (%). (G) Hepatic TG content. (H) Representative images of liver section subjected to Oil Red O staining and H&E staining (Scale bar, 100 μm). (I) Serum ALT activity, AST activity, and total-bilirubin levels. (J) qPCR assay for hepatic genes as indicated. (K) COL1A1 immunoblotting and (L) quantification for livers. (M) Representative images of liver section subjected to Sirius Red staining (Scale bar, 100 μm). Data were expressed as mean ± SEM and analyzed by Student’s t-test. * *p* < 0.05, ** *p* < 0.01.

After 7 weeks on the CDAHFD diet, control and MLKO mice (with hepatic deletion of *Mtarc1*) exhibited comparable body weights (Figure 2E). However, MLKO mice displayed significantly reduced liver-to-body weight ratios (Figure 2F) and lower hepatic triglyceride content, though cholesterol levels remained unchanged (Figure 2G, S4A). Oil Red O and H&E staining further confirmed the decrease in hepatic fat accumulation (Figure 2H). Serum markers of liver injury (ALT, AST, and total bilirubin) were consistently reduced in MLKO mice (Figure 2I), while serum triglyceride, cholesterol, and glucose levels showed no genotypic differences (Figure S4B to D). The attenuated liver injury in MLKO mice was accompanied by downregulated expression of inflammatory and fibrotic markers (Figure 2J), as well as reduced COL1A1 protein levels and diminished Sirius Red staining (Figure 2K to M), collectively suggesting improved hepatic health.

MTARC2, the closest paralog of MTARC1, shares similarities with MTARC1 in protein structure, subcellular distribution, and substrate preferences ^21, 22^. However, hepatic *Mtarc2* deletion failed to recapitulate the protective effects observed in hepatic *Mtarc1*-deficient mice (Figure S4E to J), reflecting their differential role in fatty liver disease pathogenesis. Taken together, our work demonstrated that hepatocyte-specific MTARC1 inactivation is sufficient to confer protection against diet-induced steatosis, establishing a crucial foundation for elucidating MTARC1’s molecular mechanisms in hepatic lipid metabolism.

### MTARC1 inactivation improves fatty liver disease in a lipolysis and lipophagy-dependent manner

Hepatic TG accumulation is one of the drivers of fatty liver disease. Hepatic TG homeostasis is regulated by multiple pathways, including VLDL secretion, TG degradation, and TG synthesis. To test whether MTARC1 regulates VLDL-mediated TG secretion, we employed the established poloxamer 407 inhibition method ^23^. Briefly, after overnight fasting, mice were bled, intraperitoneally injected with this lipoprotein lipase inhibitor, and re-bled. Neither global (MKO) nor liver-specific (MLKO) *Mtarc1* deletion affected VLDL-TG secretion (Figure S5A, B). In addition, inhibition of VLDL secretion did not abolish the TG-lowering effect of MTARC1 deficiency in primary hepatocytes (Figure S5C, D). ATGL-mediated lipolysis and LAL-mediated lipophagy are two major LD degradation pathways. To assess their roles in MTARC1-regulated TG homeostasis, we treated primary hepatocytes with ATGL inhibitor (ATGLi), LAL inhibitor (LALi), or both. First, individual inhibition of either pathway increased TG content (Figure 3A), supporting their roles in TG degradation. Combined inhibition had an additive effect (Figure 3A). Notably, ATGLi (but not LALi) partially attenuated the TG reduction caused by MTARC1 deficiency (Figure 3B), while dual inhibition nearly eliminated this effect (Figure 3A, B), indicating that MTARC1 deficiency reduces cellular TG content via the coordination of both TG degradation pathways. By contrast, inhibiting TG synthesis did not attenuate MTARC1 deficiency’s TG-lowering effect (Figure S5E, F).

**Fig 3.**
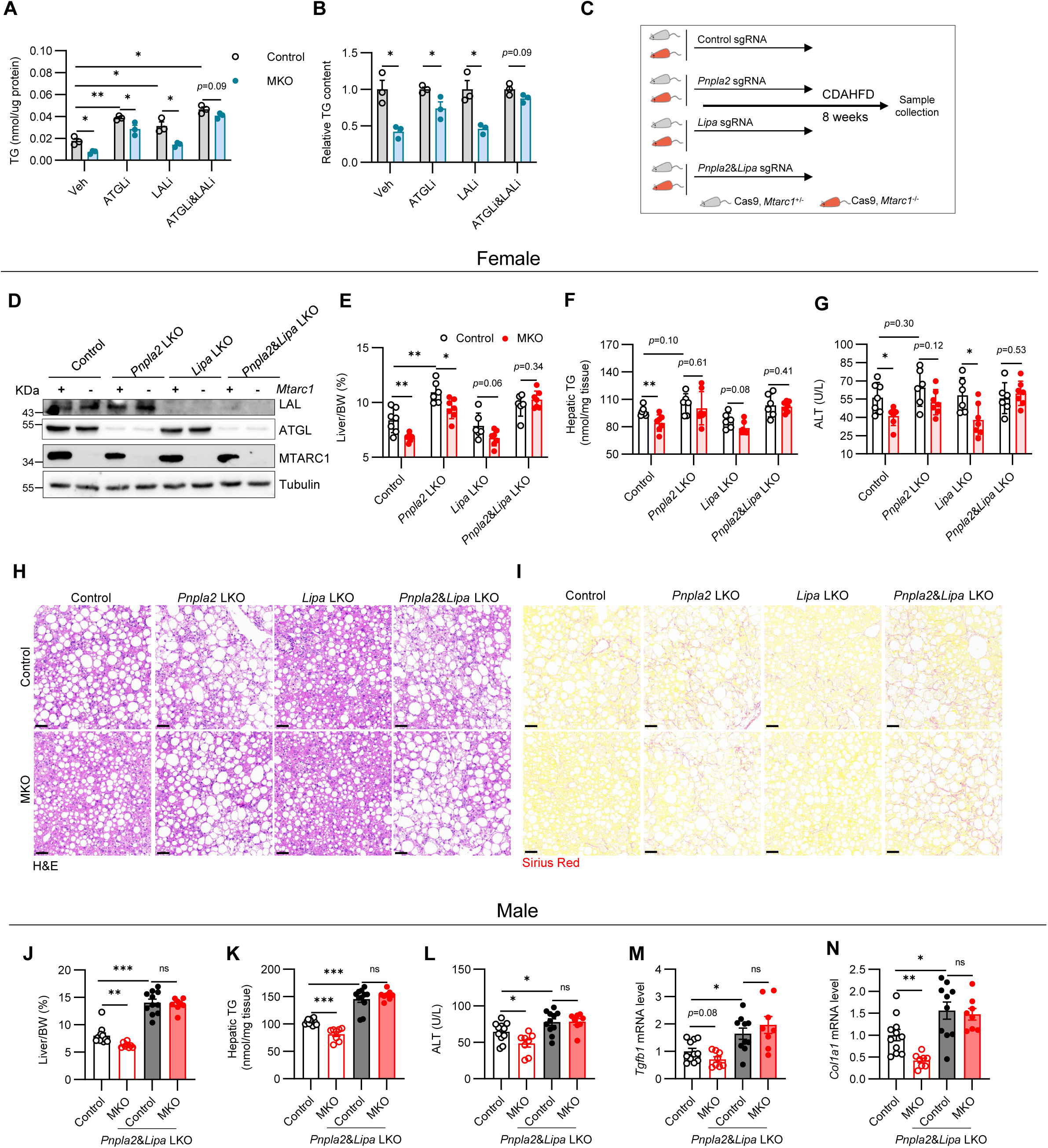
MTARC1 inactivation improves fatty liver disease in a lipolysis and lipophagy-dependent manner. (A) Primary hepatocytes were treated with the inhibitors as indicated for 24 hours and subjected to cellular TG content measurement. (B) The value of control cell was defined as one for Panel A. (C-I) Female control and MKO mice were administrated with virus as indicated and challenged with CDAHFD for eight weeks (n = 6 to 8 per group). (C) Animal experimental protocol. (D) Immunoblotting for livers against antibodies as indicated. (E) Liver body weight ratio (%). (F) Hepatic TG content. (G) Serum ALT activity. (H) Representative images of H&E staining (Scale bar, 50 μm). (I) Representative images of Sirius Red staining (Scale bar, 50 μm). (J-N) Four groups of male mouse models were generated following the protocol as shown in Panel C and sacrificed for phenotyping (n = 8 to 10 per group). (J) Liver body weight ratio (%). (K) Hepatic TG content. (L) Serum ALT activity. (M) Hepatic qPCR assay for *Tgfb1*. (N) Hepatic qPCR assay for *Col1a1*. Data were expressed as mean ± SEM and analyzed by Student’s t-test. * *p* < 0.05, ** *p* < 0.01.

To determine whether MTARC1 regulates fatty liver disease through lipolysis or lipophagy *in vivo*, we evaluated its hepatoprotective effect in mice with hepatic deletion of *Pnpla2* (encoding ATGL), *Lipa* (encoding LAL), or both (Figure 3C and D). Again, *Mtarc1* knockout alone significantly ameliorated fatty liver disease in terms of liver index, hepatic fat content, liver injury, and hepatic fibrosis (Figure 3E to I, Figure S5G), without affecting body weight, hepatic TC, serum TG, and TC (Figure S5H to K). In contrast, hepatic deletion of *Pnpla2* exacerbated fatty liver disease, whereas *Lipa* deletion had no effect (Figure 3E to I, Figure S5G), similar to the previous reports ^24, 25^. MTARC1 deficiency still attenuated fatty liver disease in *Pnpla2*- or *Lipa*-deficient livers, albeit less robustly than in controls (Figure 3E to I, Figure S5G). Strikingly, its protective effect was completely abolished in *Pnpla2* and *Lipa* double-knockout livers (Figure 3E to I, Figure S5G).

The data above were derived from female mice with a relatively small sample size. To validate these findings, we conducted an additional experiment in male mice.

Consistent with female data, dual hepatic deletion of *Pnpla2* and *Lipa* abolished the protective effect of MTARC1 deficiency (Figure 3J to N, Figure S5L, M), without altering body weight (Figure S5N). Taken together, these findings establish that the improvement in hepatic steatosis, injury and fibrosis resulting from MTARC1 deficiency requires both lipolysis and lipophagy.

### MTARC1 deficiency upregulates glycerophospholipid biosynthesis

Of note, although MTARC1 deficiency confers hepatoprotection through ATGL-mediated lipolysis and LAL-mediated lipophagy, it’s not associated with altered expression of ATGL or LAL (Figure S6A). To uncover the mechanism through which MTARC1 inactivation improves fat accumulation and liver health, we performed parallel proteomic and transcriptomic analyses of control and MLKO mouse livers (Figure 4A). Our analyses revealed distinct coverage between the two approaches: while proteomic analysis identified 5,790 proteins, transcriptomic sequencing detected expression of 15,401 genes, reflecting the greater sensitivity of mRNA detection. For downstream analysis, we integrated these datasets by considering both the 5,461 genes with corresponding protein expression data and the additional 9,940 genes identified solely at the transcript level. Differential expression analysis was then performed across this combined dataset, followed by gene ontology annotation (Figure 4A).

**Fig 4.**
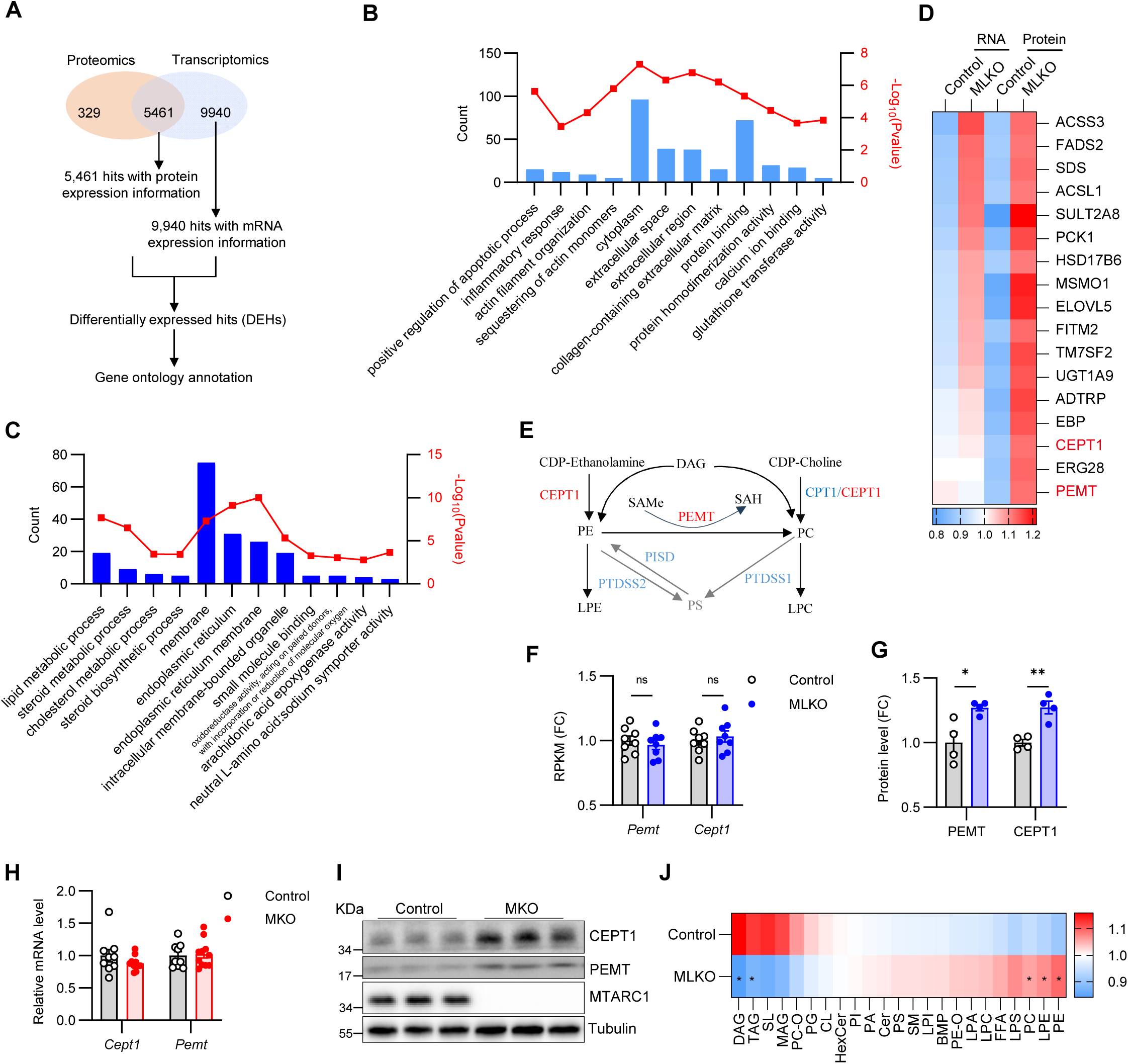
MTARC1 deficiency remodels hepatic glycerophospholipid homeostasis. (A) The workflow of proteomics and transcriptomics joint analysis. Briefly, eight livers from each group of mice challenged with CDAHFD for seven weeks were subjected to RNA-seq (n = 8/group). Two livers from the same group were pooled and subjected to TMT-based proteomics (n = 4/group). The hits with |Log_2_FC| > 0.3, *P* < 0.05 were defined as differentially expressed hits (DEHs). (B-C) Top four terms from gene ontology annotation for downregulated (B) and upregulated (C) hits (BP, biological process, CC, cellular component, MF, molecular function). (D) Heatmap presenting the detailed expression profile of the term “lipid metabolic process” from Panel C. (E) The simplified pathways through which PC and PE are synthesized. (F) RPKM level of *Pemt* and *Cept1* mined from the transcriptomic data (n = 8/group). (G) Relative protein level of PEMT and CEPT1 mined from the proteomic data (n = 4/group). (H-I) qPCR and immunoblotting assays for livers from control and MKO mice challenged with CDAHFD for eight weeks. (J) Livers from control and MLKO mice challenged with CDAHFD for seven weeks were subjected to lipidomics analysis (n = 9/group). The relative total level of each subtype identified was presented in the heatmap. Data were expressed as mean ± SEM and analyzed by Student’s t-test. * *p* < 0.05, ** *p* < 0.01.

For the downregulated hits, the top four from each category (BP, CC, and MF) were presented in Figure 4B. Of note, the inflammatory response from BP was among them, further supporting the notion that *Mtarc1* deletion suppresses inflammation (Figure 4B, Figure S6B). Another interesting term was collagen-containing extracellular matrix from CC (Figure 4B, Figure S6C), in line with the notion that *Mtarc1* deletion suppresses hepatic fibrosis. For the upregulated hits, the top four from each category (BP, CC, and MF) were presented in Figure 3C. *Mtarc1* deletion significantly decreased hepatic TG content in lipolysis and lipophagy dependent manner (Figure 3). One interesting term from BP was lipid metabolic process (Figure 4C). Within this category, two key GPL biosynthesis enzymes, CEPT1 and PEMT, showed posttranscriptional upregulation, drawing our particular interest (Figure 4D to G). Their expression profiles were further validated by western blotting and qPCR (Figure 4H and I). Of note, MTARC1 inactivation did not affect the protein stability of CEPT1 and PEMT as determined by cycloheximide chase assay (Figure S6D). These findings collectively indicate that MTARC1 modulates *Cept1* and *Pemt* expression through an unidentified post-transcriptional regulatory mechanism.

To determine whether altered CEPT1/PEMT expression modified the GPL profile, we performed comprehensive lipidomic analysis for control and MLKO livers. As anticipated, MLKO mice exhibited reduced diacylglycerol and TAG levels (Figure 4J, Table S1. Hepatic lipidomic data), while the level of total cardiolipin, a type of phospholipid essential for the normal function of mitochondria ^26^, remained unchanged (Figure 4J). No significant differences were observed in phosphatidylglycerol (PG), phosphatidylinositol (PI), alkyl PE (PE-O), phosphatidylserine (PS), ceramides (Cer), hexose ceramides (HexCer), sphingomyelin (SM), lyso-PA (LPA), bis (monoacylglycerol) phosphate (BMP), lyso-PI (LPI), and free fatty acid (FFA) (Figure 4J). By contrast, total PE and PC were significantly increased in MLKO liver, consistent with the induction of CEPT1 and PEMT (Figure 4J). Interestingly, levels of LPE were also elevated (Figure 4J), likely due to increased availability of their synthetic substrates—PE. Taken together, these data demonstrate that MTARC1 deficiency elevates *Pemt* and *Cept1* expression, thereby enhancing GPL biosynthesis.

### MTARC1 deficiency regulate lipid droplet turnover rate by remodeling glycerophospholipid

TG degradation rate is regulated by multiple factors, including LD size. As is well established, smaller LDs exhibit higher TG degradation rates due to their greater surface-to-volume ratio compared to larger LDs ^27–29^. Notably, the size of LD is regulated by both composition and abundance of GPLs ^29–31^. We observed that MTARC1 depletion elevates hepatic GPL levels by upregulating PEMT and CEPT1 (Fig 4), two transmembrane enzymes located at the ER where LDs bud off. Based on these clues, we hypothesized that in MTARC1-deficient hepatocytes, the altered GPL profile reduces the size of LDs, thereby increasing their surface-to-volume ratio and enhancing LD degradation through lipophagy and lipolysis, which ultimately reduces cellular TG content.

To address the hypothesis above, we isolated LDs from livers of wild-type C57 mice (Figure 5A) and determined their GPL composition by lipidomics (Figure 5B). As expected, PC (67.5%) and PE (13.6%) were the two most abundant GPL subtypes in hepatic LDs (Figure 5B). LPE, the third most abundant GPL subtype, was present at levels similar to PE (Figure 5B). Interestingly, the major GPL of hepatic LDs were those upregulated by MTARC1 deficiency (Figure 5C). LPE, a type of lysolipids, could be synthesized via phospholipid hydrolysis carried out by the phospholipase A2 (PLA2) superfamily (Figure 5D)^32^. Su *et al* reported that PLA2G4C, one enzyme from the PLA2 family, could colocalizes with LDs ^33^. Consistent with this report, mCherry-PLA2G4C (but not mCherry alone) colocalized with LDs (Figure 5E), suggesting that LD-associated LPE may be locally synthesized. Taken together, the GPLs upregulated upon MTARC1 depletion represent major components of hepatic LDs.

**Fig 5.**
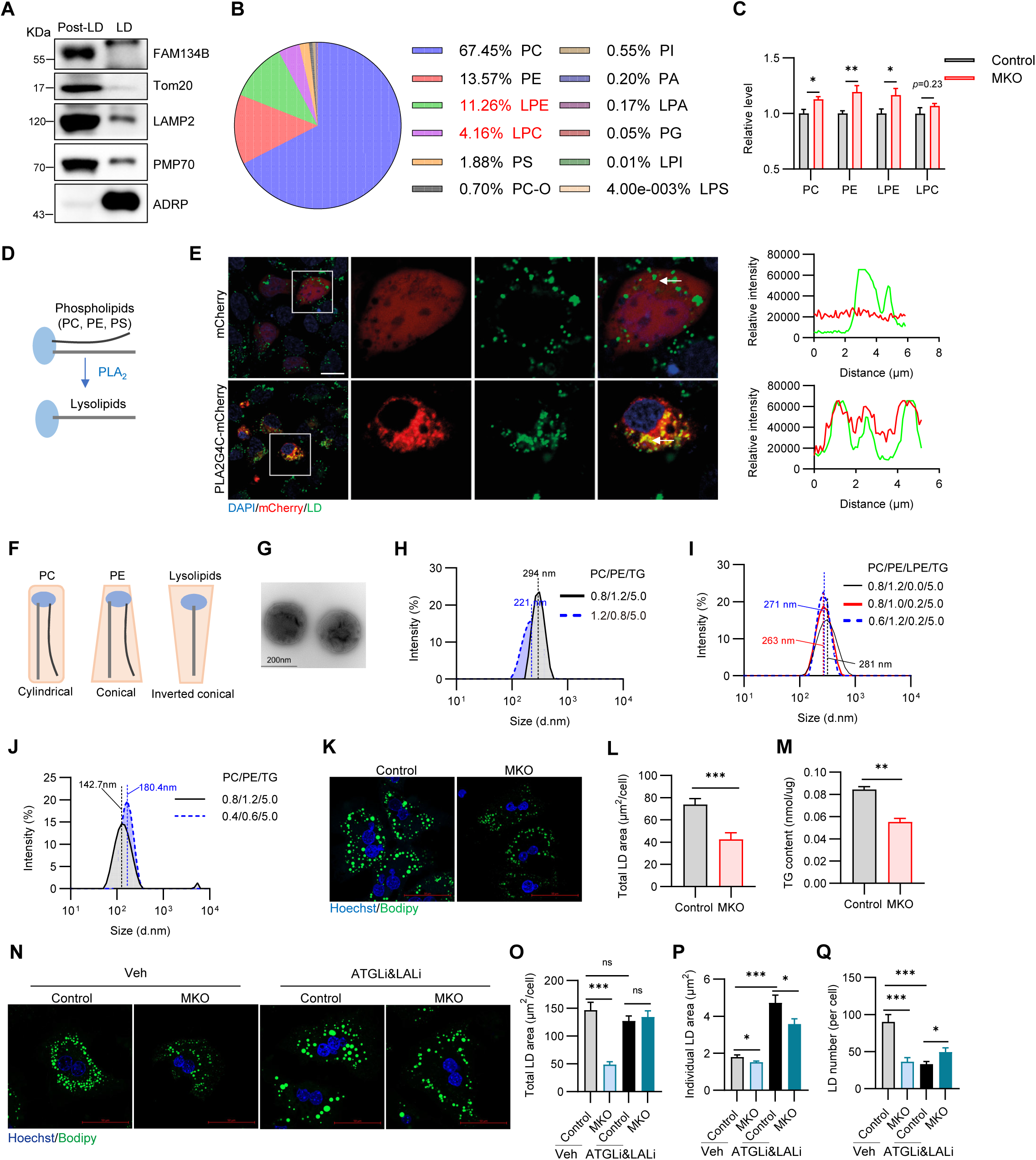
MTARC1 deficiency regulate lipid droplet turnover rate by remodeling glycerophospholipid. (A) The LD and post-LD fractions from mouse livers were subjected to immunoblotting against antibodies as indicated. FAM134B, ER marker; Tom20, mitochondria marker; LAMP2, lysosome marker; PMP70, peroxisome marker; ADRP, LD marker. (B) Pie chart presenting the phospholipid compositions of hepatic LDs. (C) Total hepatic PC, PE, LPE, and LPC mined from hepatic lipidomic data related to Fig. 4J (n = 9/group). (D) The simplified pathway through which lysolipids were synthesized. (E) HEK293T cells were transfected with the plasmids either expressing mCherry alone or PLA2G4C-mCherry fused protein. The cells were subjected to BODIPY staining and imaging after 24 hours post-transfection (Scale bar, 20 μm). (F) Geometry of PC, PE, and lysolipids. (G) A representative image of artificial adiposome captured by TEM (Scale bar, 200 nm). (H-J) Artificial adiposome assay; PC, DOPC; PE, DOPE; LPE, LPE18:0; and TG, Triolein (unit, mg). (H) The increase of PC/PE ratio reduces the size of adiposome. (I) The substitution of LPE for PC or PE decreases the size of adiposome. (J) The increase of PC and PE amount reduces the size of adiposome. (K-M) Primary hepatocytes were isolated from control and MKO mice and cultured for 36 hours, and subjected to Hoechst and Bodipy staining. (K) Representative images of cells subjected to Bodipy staining (Scale bar, 50 μm). (L) Total LD area per cell (n = 20/group). (M) TG content in primary hepatocyte from control and MKO mice (n = 3/group). (N-Q) Primary hepatocytes were isolated from control and *Mtarc1* knockout mice and subjected to downstream analyses. (N) The cells were cultured with either vehicle or ATGLi&LALi for 24 hours and subjected to Bodipy staining and imaging. (O) Total cellular LD area per cell (n = 28-41 cells/group). (P) Individual LD area (n = 1261-3063 LDs/group). (Q) LD number per cell (n = 28-41 cells/group). Data were expressed as mean ± SEM and analyzed by Student’s t-test. * *p* < 0.05, ** *p* < 0.01.

Based on the geometry of GPLs (Figure 5F), we reasoned that LPE has a stronger LD size-reducing effect than both PC and PE. To test this, we generated artificial adiposome and confirmed their formation by TEM (Figure 5G). Higher ratios of PC/PE had been reported to reduce adiposome size ^34^, which we confirmed in our system (Figure 5H), validating our experimental approach. LPE18:0 was the most abundant LPE species in LDs (Table S2. Hepatic LD LPE composition). As expected, substituting LPE18:0 for an equal amount of PE or PC reduced artificial adiposome size (Figure 5I). Furthermore, GPL amount was inversely correlated with adiposome size (Figure 5J), consistent with reports that GPL levels negatively regulate LD size ^29–31^. These data suggests that remodeled GPL induced by MTARC1 deficiency might have a LD size-reducing effect. In line with this possibility, BODIPY staining of primary hepatocytes revealed smaller mean LD size in MKO cells (Figure 5K, L). We also observed reduced TG content in hepatocytes lacking MTARC1 (Figure 5N), making it difficult to determine the primary effect. Because dual inhibition blunted the TG-lowering effect of MTARC1 deficiency, we examined LD morphology under this condition (Figure 5N to Q). Remarkably, under dual inhibition, MKO cells exhibited smaller but more numerous LDs compared to controls (Figure 5P, Q), supporting the hypothesis that MTARC1 deficiency enhances GPL levels, which in turn reduce LD size. Collectively, these data suggest that MTARC1 deficiency elevates hepatic GPLs, reducing LD size and increasing surface-to-volume ratio. This might in turn enhances LD degradation through lipolysis and lipophagy, ultimately decreasing cellular TG content.

### Inhibition of glycerophospholipid biosynthesis reverses the phenotype of MTARC1 null mice

While MTARC1 deficiency effectively ameliorated CDAHFD-induced fatty liver disease (Figure 1), it showed no protective effect against HFHFD-induced steatosis (Figure S3). This dietary specificity may relate to choline deficiency in CDAHFD, as choline serves as the essential substrate for GPL biosynthesis. These data highlight the role of enhanced GPL biosynthesis in the improvement of hepatic steatosis. To directly test PEMT-dependency, we examined MTARC1 deficiency in hepatocyte-specific *Pemt* knockout mice (Figure 6A, Figure S7A). As previously reported ^35^, hepatic *Pemt*-deficient mice on choline-deficient diet exhibited severe body weight loss (Figure 6B), necessitating experiment termination at 3 weeks. At this study endpoint, MKO mice exhibited a nonsignificant reduction in liver-to-body weight ratio (Figure S7B). However, MLKO mice demonstrated significantly reduced ALT levels compared to controls, a protective effect that was substantially diminished when *Pemt* was concomitantly deleted (Figure 6C). Intriguingly, hepatocyte-specific *Pemt* deletion alone led to increased liver body weight ratio without corresponding changes in hepatic TG content (Figure S7B and C), likely, complete ablation of hepatic *Pemt* have pleiotropic effects on cellular TG metabolism.

**Fig 6.**
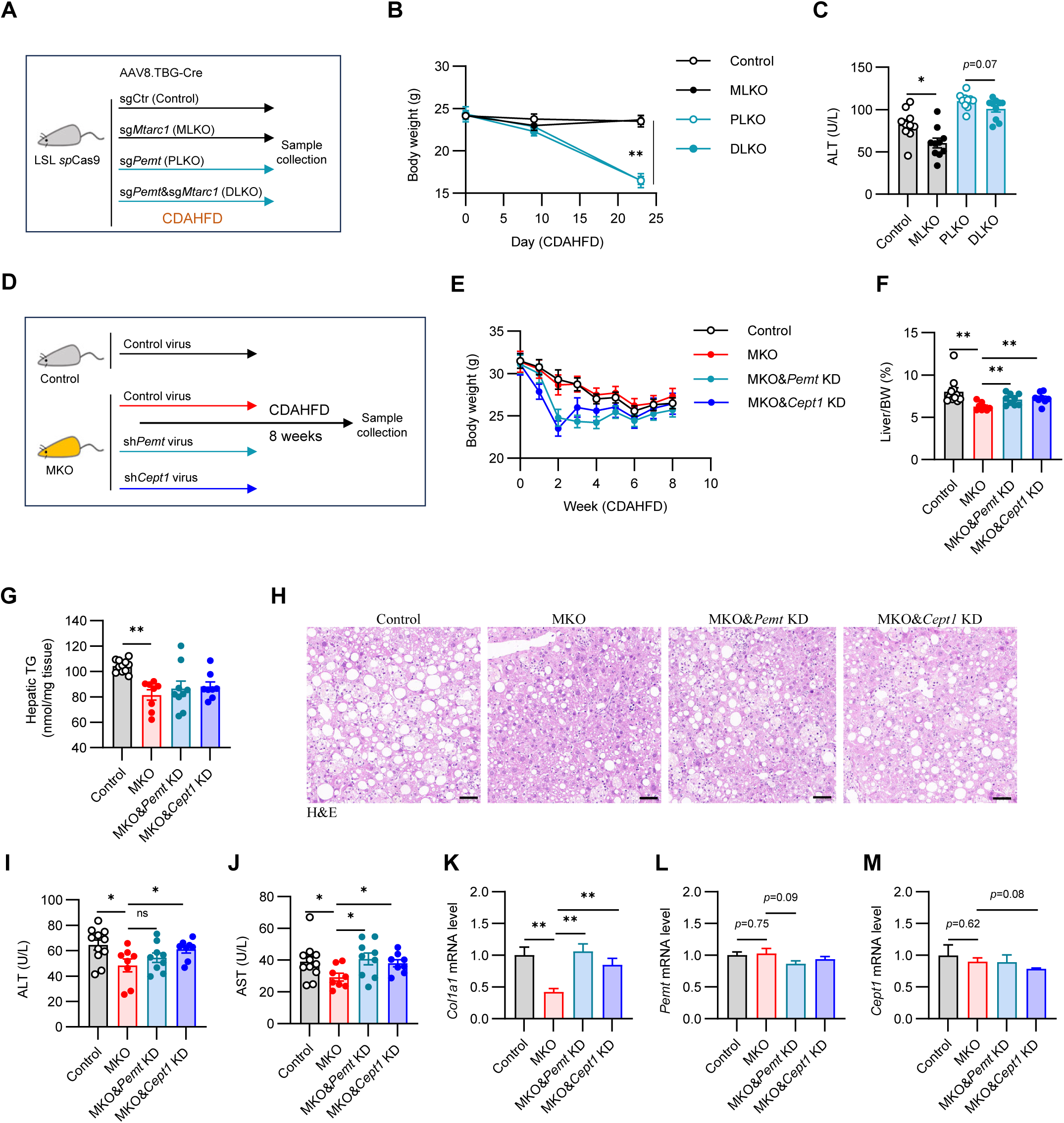
Inhibition of glycerophospholipids biosynthesis reverses the hepatoprotective effect of MTARC1 deficiency. (A-C) Eight weeks old LSL Cas9 mice were administrated with the virus as indicated and subjected to CDAHFD challenge for 23 days (n = 9-10/group). The viral dosages (vg/mouse) are: 2e^10^ for *Pemt* shRNA virus, 3e^10^ for *Cept1* shRNA, and 1e^11^ for control virus. To ensure the equal amount of virus used for each mouse, control virus was used as the filler. (A) Animal diet challenge experiment workflow. (B) Growth curve. (C) Serum ALT activity. (D-M) Eight weeks old control and *Mtarc1* knockout mice were administrated with virus as indicated and subjected to CDAHFD challenge for eight weeks (n = 9-10/group). (D) Animal experimental protocol. (E) Growth curve. (F) Liver body weight ratio (%). (G) Hepatic TG content. (H) Representative images of H&E staining for liver section (Scale bar, 50 μm). (I) Serum ALT activity. (J) Serum AST activity. (K-M) qPCR assay for Hepatic *Col1a1* (K), *Pemt* (L), and *Cept1* (M). Data were expressed as mean ± SEM and analyzed by Student’s t-test. * *p* < 0.05, ** *p* < 0.01.

Next, we adopted a partial knockdown strategy to more precisely evaluate the roles of *Pemt* and *Cept1* upregulation in the MKO phenotype (Figure 6D). MKO mice were administered with AAV8 vectors expressing sh*Pemt* or sh*Cept1* at titers achieving partial knockdown (Figure S7D, E). Partial *Pemt*/*Cept1* inhibition initially reduced body weights, but this effect resolved within three weeks (Figure 6E). After a 6-week model diet challenge, VLDL-mediated TG secretion was performed. Surprisingly, neither *Cept1* nor *Pemt* knockdown affected the TG secretion rate (Figure S7F), likely because the partial knockdown was too modest to produce a measurable impact. However, the liver body weight ratio of MKO mice reverted to control levels with either sh*Pemt* or sh*Cept1* treatment at 8 weeks (Figure 6F). While hepatic TG content exhibited an increasing trend that did not reach statistical significance (Figure 6G and H), TC levels remained comparable across all interventions (Figure S7G). Liver injury markers (ALT/AST) and *Col1a1* expression mirrored the liver body weight ratio patterns (Figure 6I to K). Notably, residual *Pemt*/*Cept1* knockdown persisted at termination (*p*=0.09 and *p*=0.08, respectively; Figure 6L, M), with *Mtarc1* deletion confirmed by qPCR (Figure S7H). These findings establish that PEMT/CEPT1 upregulation mediates a substantial portion, though likely not all, of MTARC1 deficiency’s hepatoprotective effects.

## Discussion

Human genetic studies have identified MTARC1 variants linked to metabolic dysfunction-associated fatty liver disease (MASLD), suggesting its therapeutic potential. While MTARC1’s role in preclinical MASLD models remains elusive, our work establishes it as a key disease driver. Strikingly, global MTARC1 inactivation significantly attenuated diet-induced hepatic fat accumulation, liver injury, inflammation, and fibrosis, in both males and females. Liver-specific knockout recapitulated these protective effects. Furthermore, we demonstrated that MTARC1 inactivation’s hepatoprotection requires both lipolysis and lipophagy. Through multi-omics integration (transcriptomics, proteomics, and lipidomics), we revealed that MTARC1 deficiency elevates GPL levels (PC, PE, and LPE) through post-transcriptional upregulation of CEPT1 and PEMT, reducing LD size while increasing their surface-to-volume ratio, which partially explains how MTARC1 deficiency improves fatty liver disease via coordinated lipolysis and lipophagy (Figure 7).

**Fig 7.**
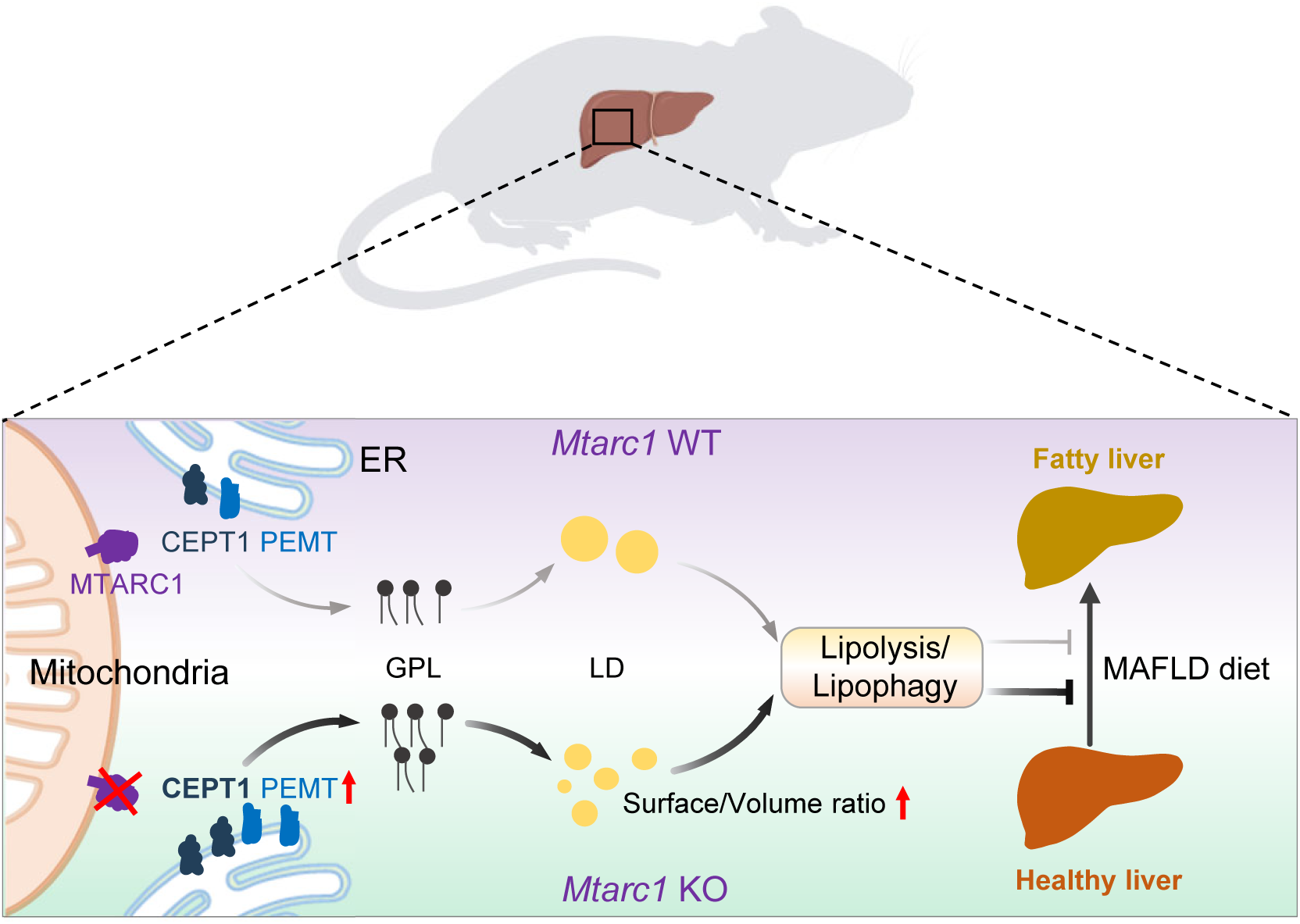
A model depicting the proposed molecular mechanism through which MTARC1 regulates the progression of fatty liver disease.

Human genetic analyses have linked MTARC1 to MASLD, while no such link has been found for MTARC2, a MTARC1’s closest paralog. Nevertheless, emerging evidence suggests that MTARC2 may also participate in lipid metabolism. This is supported by two *in vitro* studies in 3T3-L1 adipocytes demonstrating that *Mtarc2* knockdown reduces cellular TG content ^36, 37^. In addition, MTARC2 shares similar protein structure, subcellular localization, and partially overlapping substrate preferences ^21, 22^. These observations suggest that MTARC1 and MTARC2 may play analogous roles in hepatic lipid metabolism. However, hepatic MTARC2 deficiency failed to recapitulate the hepatoprotective effects observed with MTARC1 knockout, underscoring their differential involvement in fatty liver disease pathogenesis.

LDs are highly dynamic organelles that frequently interact with the ER, peroxisomes, and mitochondria ^30^. In hepatocytes, TG stores are mobilized through two principal pathways, lipolysis and lipophagy, with the liberated fatty acids subsequently undergoing β-oxidation in peroxisomes and/or mitochondria. The tight coupling between TG mobilization and β-oxidation is well established, with substrate availability serving as the primary driver of metabolic flux ^14, 17, 38, 39^. This mechanistic understanding positions TG mobilization pathways as promising therapeutic targets for reducing hepatic lipid accumulation. Interestingly, a study by Ester *et al* demonstrated that MTARC1 knockdown enhances fatty acid β-oxidation in human primary hepatocytes ^40^, an effect likely attributable to increased fatty acid substrate availability. Dysregulated phospholipid homeostasis is implicated in MASLD pathogenesis across preclinical models and clinical observations. The deletion of *Pemt* in mice leads to liver failure, and fatty liver ^35, 41, 42^. These findings are clinically relevant, as PEMT polymorphisms correlate with human MASLD susceptibility ^43^. In addition, hepatic deficiency of CEPT1, an enzyme involved in both PC and PE synthesis via the CDP pathway, leads to fatty liver as well ^44^. Individual inhibition of CEPT1 or PEMT completely rescues the effect of MTARC1 deficiency on liver body weight ratio, liver injury and liver fibrosis, but only partially normalizes hepatic TG levels (Fig 6). Likely, combined inhibition of CEPT1 and PEMT are necessary to normalize the altered GPL synthetic activity. Of note, the precise molecular mechanisms through which MTARC1 regulates the expression of CEPT1 and PEMT remain to be elucidated and warrant further investigation.

We proposed the mechanistic model that by upregulating PEMT and CEPT1, MTARC1 deficiency increases the levels of PC, PE, and LPE—three major LD GPLs, enhancing LD degradation through lipolysis and lipophagy by reducing LD size and increasing their surface-to-volume ratio. Both LDs and VLDLs bud from the ER, where PEMT and CEPT1 synthesize GPLs ^30^. While the mechanisms governing their differential sorting remain unclear, bud-associated protein composition appears to be a determining factor ^30^. Notably, MTARC1 deficiency upregulates FITM2 (Figure 4D), a well-characterized pro-LD factor ^45–47^. Of note, GPL alterations exert distinct metabolic effects depending on their subcellular localization ^48^. CDP-choline pathway-mediated PC biosynthesis could be carried out through two spatially distinct enzymes (Figure 4E): ER-localized CEPT1 and Golgi-resident CPT1 ^18^. Intriguingly, MTARC1 deficiency selectively upregulates CEPT1 (ER) but not CPT1 (Golgi), revealing a compartment-specific regulation of GPL synthesis. Likely, these two pieces of fact might explain why MTARC1 deficiency ameliorates fatty liver through GPL-dependent mechanisms that preferentially enhance LD degradation without affecting VLDL-associated TG secretion.

In summary, our work establishes MTARC1 as a critical regulator of hepatic lipid homeostasis. MTARC1 constrains GPL biosynthesis by modulating CEPT1 and PEMT through an undefined mechanism, thereby influencing LD-TG mobilization and liver metabolic health. Targeting MTARC1 has the potential to be an effective strategy for treating metabolism-associated fatty liver disease.

## Materials and Methods

### Mouse models and animal experiments

Animal experiments were performed following the guidelines provided by the Ethics Committees of Anhui Medical University. All mice had free access to water and food, and were housed in a specific pathogen-free environment with a 12-h light/12-h dark cycle. *Mtarc1* knockout mice on C57BL/6J genetic background were generated using the CRISPR/Cas9 system by GemPharmatech (Nanjing, China). Rosa26-CAG-LSL-Cas9 mice on C57BL/6J genetic background were also obtained from GemPharmatech (Nanjing, China). The mice were fed either standard rodent chow, a diet composed of 62% kcal fat, choline-deficient, with 0.1% methionine (CDAHFD) (Research diets, A06071302), or 40% kcal fat, 20% kcal fructose and 2% cholesterol (HFHFD) (Research diet, D09100310) as indicated. The primer sets used for *Mtarc1* null mice genotyping are as follows: *F1-atg tat tcc tgc atg cca gaa gag*, and *R1-ccc tac atc act acg cat gtt cc*; *F2-agg gtc tga acg aat aga ttc cca*, and *R2-cat cca aag gtt gtc aca aag gta ag*. The binding sites of these primers were presented in Figure S1A. LSL-Cas9 mice were tail vein injected with AAV8 to generate mouse models as indicated in figure legends at the dosage of 1e^11^ to 5e^11^ vg/mouse unless stated otherwise. The first two groups were shared between Figure 3J-K and Figure 6D-M. The mice were fed either standard rodent chow, a diet composed of 62% kcal fat, choline-deficient, with 0.1% methionine (CDAHFD) (Research diets, A06071302), or 40% kcal fat, 20% kcal fructose and 2% cholesterol (HFHFD) (Research diet, D09100310) as indicated.

### Cell lines

HEK293T cells and HepG2 were maintained at 37 ^0^C and 5% CO_2_ and cultured in Dulbecco’s modified Eagle’s medium (Gibco) supplemented with 10% fetal bovine serum (HyClone) and 100 IU penicillin and 100 mg/mL streptomycin (P/S).

### Primary hepatocytes

Mouse primary hepatocytes were isolated by collagenase perfusion as described previously ^15^. Briefly, collagenase digestion was performed with portal vein perfusion and physical dissociation of digested tissue. Hepatocytes were isolated by multiple centrifugation steps, including a Percoll (Biosharp, BS909) gradient for separation of viable cells, subsequently plated onto collagen-coated coverslips or culture dishes, and were maintained in Dulbecco’s modified Eagle’s medium. After overnight incubation, hepatocytes were used for downstream experiments. For Lomitapide treatment, 1 µM Lomitapide (MCE, HY-14667) was used to inhibit VLDL assembly for 24 h. Both 10 µM DGAT1 inhibitor PF-04620110 (MCE, HY-13009) and 10 µM DGAT2 inhibitor PF-06424439 (MCE, HY-108341) were added to inhibit TG synthesis after 12 h of incubation with 50 µM oleic acid. Atglistatin (MCE, HY-15859, 50 µM) and Lalistat 1 (MCE, HY-116815, 50 µM) were used to inhibit lipolysis and lipophagy, respectively.

### Immunofluorescence

Primary hepatocytes cultured on coverslips were fixed with 4% paraformaldehyde, followed by incubation with BODIPY 493/503 (Beyotime, C2053S) for neutral lipid staining. Nuclei were stained with Hoechst (Beyotime, P0133). Images were acquired with a Laser Confocal Microscope (Carl Zeiss, LSM980) with a 63× magnification.

HEK293T cells were transfected with pLJM1-CMV-mCherry (control) or pLJM1-CMV-mCherry-PLA2G4C plasmids, followed by BODIPY staining and fluorescence imaging to evaluate lipid droplet. All images were analyzed using Zen Blue (Carl Zeiss) and ImageJ (NIH).

### Serum biochemical assay

Serum TG (Jiancheng, A110-1-1), total cholesterol (Jiancheng, A111-1-1), glucose (Jiancheng, A154-1-1), non-esterified fatty acid (NEFA) (Jiancheng, A042-2-1), alanine aminotransferase (ALT) (Jiancheng, C009-2-1), aspartate aminotransferase (AST) (Jiancheng, C010-2-1), and total-bilirubin levels (SolarBio, BC5185) were measured according to the instructions provided by the manufacturers, with some assay performed using an automated biochemical analyzer (Mindary, China).

### Tissue biochemical assay

Liver lipids were extracted for TG and cholesterol quantification using the same kits as for serum measurements. Briefly, tissues were homogenized in 15 volumes of sterilized water and 32 volumes of chloroform/methanol (2/1), followed by centrifugation at 1200g, room temperature, for 10 minutes. The chloroform phase was collected, evaporated for TG and cholesterol analysis.

### Triglyceride secretion assay

The TG secretion rate was measured as previously described ^15^. Briefly, mice were subjected to overnight fast, bled, and then IP injected with 1g/kg of poloxamer 407. Blood was collected again 5 hours post-injection, and serum TG levels were measured to calculate the secretion rate.

### Adeno-associated virus production

For AAV.TBG.Cre.U6.sgRNA.U6.sgRNA plasmids, the construction was performed as previously described ^49^. To construct AAV.U6.shRNA.U6.shRNA plasmids, AAV.TBG.Cre (Addgene #107787) was first used to generate AAV.EcoRI.U6.AgeI.Filler.HindIII, which was named pAAV.U6.shFiller. Next, the shRNA of a gene of interest was cloned into pAAV.U6.shFiller linearized with *Age*I and *Hind*III using a standard protocol. Then, U6.shRNA fragments were amplified by PCR using pAAV.U6.shRNA as a template and cloned into pAAV.U6.shFiller linearized with *Eco*RI and *Hind*III using an in-fusion cloning strategy. sgRNA and shRNA sequences were provided in Table S3. List of sgRNA and shRNA sequence. The AAV8 packaging and purification were performed as previously described ^49^.

### Hepatic transcriptomics and proteomics

Transcriptomic and proteomic analyses were conducted as previously described ^50^. Briefly, liver samples were harvested and submitted to Seqhealth Technology Co., LTD (Wuhan, China) for the transcriptomic analysis based on the Illumina Seq method. Raw reads of RNA-seq were deposited to the National Center for Biotechnology Information’s Sequence Read Archive under accession no. PRJNA1158946. The proteomic analysis using isobaric labeling through tandem mass tag (TMT) was performed by Oebiotech Company (Shanghai, China). The proteomic data were deposited to the iProx database with accession number PXD044540.

### Hepatic lipidomics

Liver tissues were subjected to lipids extraction using a modified version of Bligh and Dyer’s method as described previously ^51^. Briefly, tissues were homogenized in 900 µL of chloroform: methanol: MilliQ H_2_O (3:6:1) (v/v/v). The homogenate was then incubated at 1500 rpm for 30min at 4℃. At the end of the incubation, 350 µL of distilled water and 300 µL of chloroform were added to induce phase separation. The samples were then centrifuged and the lower organic phase containing lipids was extracted into a clean tube. The lipid extracts were dried in the SpeedVac under OH mode. Samples were stored at −80 ℃ until further analysis.

Lipidomic analyses were conducted by LipidALL Technologies using a Jasper HPLC coupled with Sciex TRIPLE QUAD 4500 MD as reported previously ^52–54^. Separation of individual lipid classes by normal phase (NP)-HPLC was carried out using a TUP-HB silica column (i.d. 150×2.1 mm, 3 µm) under the following conditions: mobile phase A (chloroform: methanol: ammonium hydroxide, 89.5:10:0.5) and mobile phase B (chloroform: methanol: ammonium hydroxide: water, 55:39:0.5:5.5). MRM transitions were set up for comparative analysis of various lipids. Individual lipid species were quantified by referencing to spiked internal standards. d9-PC32:0(16:0/16:0), PE 34:0, dic8-PI, d31-PS, C17:0-PA, DMPG, CL-14:0, C14-BMP, C12-SL, C17-LPC, C17:1-LPI, C17:0-LPA, C17:1-LPS, C17-Cer, C12-SM, C8-GluCer, C8-LacCer, Gb3-d18:1/17:0, d3-16:0 carnitine, and DAG(18:1/18:1)-d5 were obtained from Avanti Polar Lipids. GM3-d18:1/18:0-d3 was purchased from Matreya LLC. Free fatty acids were quantitated using d31-16:0 (Sigma-Aldrich). d6-CE 18:0 and TAG (16:0) 3-d5 were obtained from CDN isotopes.

### Hepatic Lipid droplet isolation

Hepatic LDs were isolated as previously described ^55^. Briefly, the fresh mouse liver tissue was homogenized in buffer A (250 mM sucrose, 20 mM glycine, 0.2 mM PMSF, pH 7.8 adjusted with potassium hydroxide) at a ratio of 1:3 (w/v), followed by centrifugation at 100 g for 10 min at 4°C. The supernatant was centrifuged at 3000 g for 30 minutes at 4°C. The upper layer was the LD fraction, and the precipitate is post-LD fraction. Both fractions were rinsed with buffer B (20 mM HEPES, 100 mM KCl, 2 mM MgCl_2_) at a ratio of 1:1. Both of them were either immediately subjected to downstream experiments or stored at −80°C for late use.

### Histology analysis

Oil Red O staining was performed as previously described ^56^. Briefly, tissues were harvested and transferred to a 4% paraformaldehyde solution overnight. The tissue was subsequently incubated overnight in a 30% sucrose solution before being embedded in OCT compound and stored at −20°C overnight and then sectioned. Slides were then dipped in 10% formalin for 10 minutes, 60% isopropanol once, Oil Red O solution for 15 minutes, 60% isopropanol once, and then distilled water 10 times before mounting coverslips and imaging.

H&E staining was performed with a standard protocol. Briefly, paraffin sections were subjected to dewax and hydration before staining. The sections stained with hematoxylin and eosin were sealed with neutral gum before imaging. Sirius Red staining was performed as reported ^57^. Briefly, the dewaxed and hydrated paraffin sections were stained in a Sirius Red staining solution for 8 minutes and then dehydrated quickly in three cylinders of anhydrous ethanol. The slides were subsequently dipped in the xylene for 5 minutes and sealed with neutral gum for imaging. The quantification was conducted using ImageJ.

### Plasmid construction

mCherry fragment was prepared by PCR using the plasmid pLenti-myc-GLUT4-mCherry (Addgene: #64049) as a template and cloned into pLJM1-GFP (Addgene #19319) digested with *Age*I and *Eco*RI, to generate the plasmid pLJM1-CMV-mCherry. hPLA2G4C was PCR-amplified from HepG2 cDNA and cloned into *Eco*RI-linearized pLJM1-CMV-mCherry to construct pLJM1-CMV-mCherry-PLA2G4C. The in-fusion strategy was employed to clone the plasmid above following the instructions provided in the kit (Vazyme, C115).

### Immunoblotting

Tissues were lysed in RIPA lysis buffer supplemented with 1 mM phenylmethanesulfonylfluoride and protease inhibitor cocktail (Biosharp, #BL612A) as described previously ^58^. Total protein concentrations were measured by BCA kit (Biosharp, #BL521A). The supernatant was mixed with SDS loading buffer containing β-mercaptoethanol and boiled at 95℃ for 10 minutes. An equal amount of total protein for each sample was submitted to run SDS-PAGE with a standard protocol. Primary and secondary antibodies used in this study are listed below: MTARC1 (Aviva, ARP49748_P050), HSP60 (Proteintech, 15282-1-AP), COL1A1 (Zenbio, 501352), CEPT1 (Proteintech, 20496-1-AP), PEMT (Invitrogen, PA5-42383), Tubulin (Beyotime, AF0001), LAL (Abcam, ab154356), ATGL (Proteintech, 55190-1-AP), FAM134B (Proteintech, 21537-1-AP), Tom20 (Proteintech, 11802-1-AP), LAMP2 (Proteintech, 11802-1-AP), PMP70 (Abcam, ab85550), and ADRP (Zenbio, R381796).

### Quantitative real-time PCR (qPCR)

Quantitative real-time PCR (qPCR) was performed as previously described ^58^. Total RNA was extracted using Total RNA Extraction Reagent (Vazyme, #R401) and reverse transcribed using Hifair® Ⅲ 1st Strand cDNA Synthesis SuperMix for qPCR (gDNA digester plus) (Yeasen, #11141ES60). qPCR was performed using ChamQ Universal SYBR qPCR Master Mix (Vazyme, #Q711) according to the instructions provided in the kit. qPCR primers targeting sgRNA cleavage sites were designed for knockout efficiency validation. The primers used were listed in Table S4. List of qPCR Primers.

### Artificial adiposome assay

The artificial adiposome assay was performed as previously described ^34^. Briefly, 2 mg of phospholipids (DOPC, P6354, sigma; DOPE, 850725C, Sigma; lyso-PE, 856715P, Sigma) were mixed in a 1.5ml centrifuge tube at the indicated mass ratios. The mixture was dried under nitrogen gas flow after bringing the volume to 100 μL with chloroform. 100 μL of prewarmed buffer B (20 mM HEPES, 100 mM KCl, 2 mM MgCl_2_, pH 7.4) was added at 37 °C, followed by 5 mg triolein (T7140, Sigma). The mixture was vortexed (10 seconds on and 10 seconds off, 24 cycles) until milky white, then centrifuged at 4 °C, 20,000 g for 5 minutes. The floating white band (containing adiposomes) was collected, resuspended in 100 μL buffer B, and centrifuged again (20,000 g for 5 minutes).

The floating white band was collected and resuspended in 100 μL buffer B, and centrifuged at 1000 g for 5 minutes. After removing large aggregates on the top, purified adiposomes were stored at 4 °C for subsequent analysis. Adiposome size was determined by dynamic light scattering (Zetasizer Ultra, Malvern). For TEM, samples were adsorbed onto carbon film-coated grids (10 minutes), stained with 10 μL of 0.1% tannic acid (5 minutes) and 2% uranyl acetate (5 minutes), rinsed three times with double-distilled water, and imaged using an HT7800 microscope (Hitachi).

### Statistics

Graphing was performed using GraphPad Prism 9. Experimental data were expressed as the mean ± SEM. Statistical significance between groups was performed using the student’s t-test provided in Excel, with *P* values of<0.05 were defined as statistical significance.

## CRediT authorship contribution statement

**Meng Tie**, **Liwei Hu**, **Yunzhi Yang**: Investigation, Data curation, Formal analysis. **Shaoxuan Song**, **Qihan Zhu, Jun Li, Wenjing Wang, Peng Xu, Juan Yu, Mengyue Wu, Tianheng Zhao, Delong Yuan**: Investigation. **Hongyu Bao**, **Xiuyun Wang**: Methodology. **Irfan J. Lodhi**, review & editing. **Yong Chen**: Supervision, Writing – review & editing. **Yali Chen**: Supervision, Writing – review & editing. **Anyuan He**: Conceptualization, Data curation, Formal analysis, Investigation, Methodology, Supervision, Validation, Visualization, Writing – original draft, Writing – review & editing.

## Data availability

The RNA sequencing data have been deposited in the National Center for Biotechnology Information’s Sequence Read Archive under accession no. PRJNA1158946. The proteomic data have been deposited in the iProx database under accession no. PXD044540. The data that support the findings of this study are available on request from the corresponding author.

## ABBREVIATIONS

ALT: alanine aminotransferase
AST: aspartate aminotransferase
CEPT1: choline/ethanolamine phosphotransferase 1
GPL: glycerophospholipid
LD: lipid droplet
MASLD: metabolic dysfunction-associated steatotic liver disease
MKO: *Mtarc1* knockout
PC: phosphatidylcholine
PE: phosphatidylethanolamine
PEMT: phosphatidylethanolamine N-methyltransferase
TG: triglyceride
VLDL: very low-density lipoprotein.

