## Supplemental Figures for "MTARC1 Regulates Lipid Droplet Degradation via Phospholipid Remodeling in Metabolic Fatty Liver Disease"

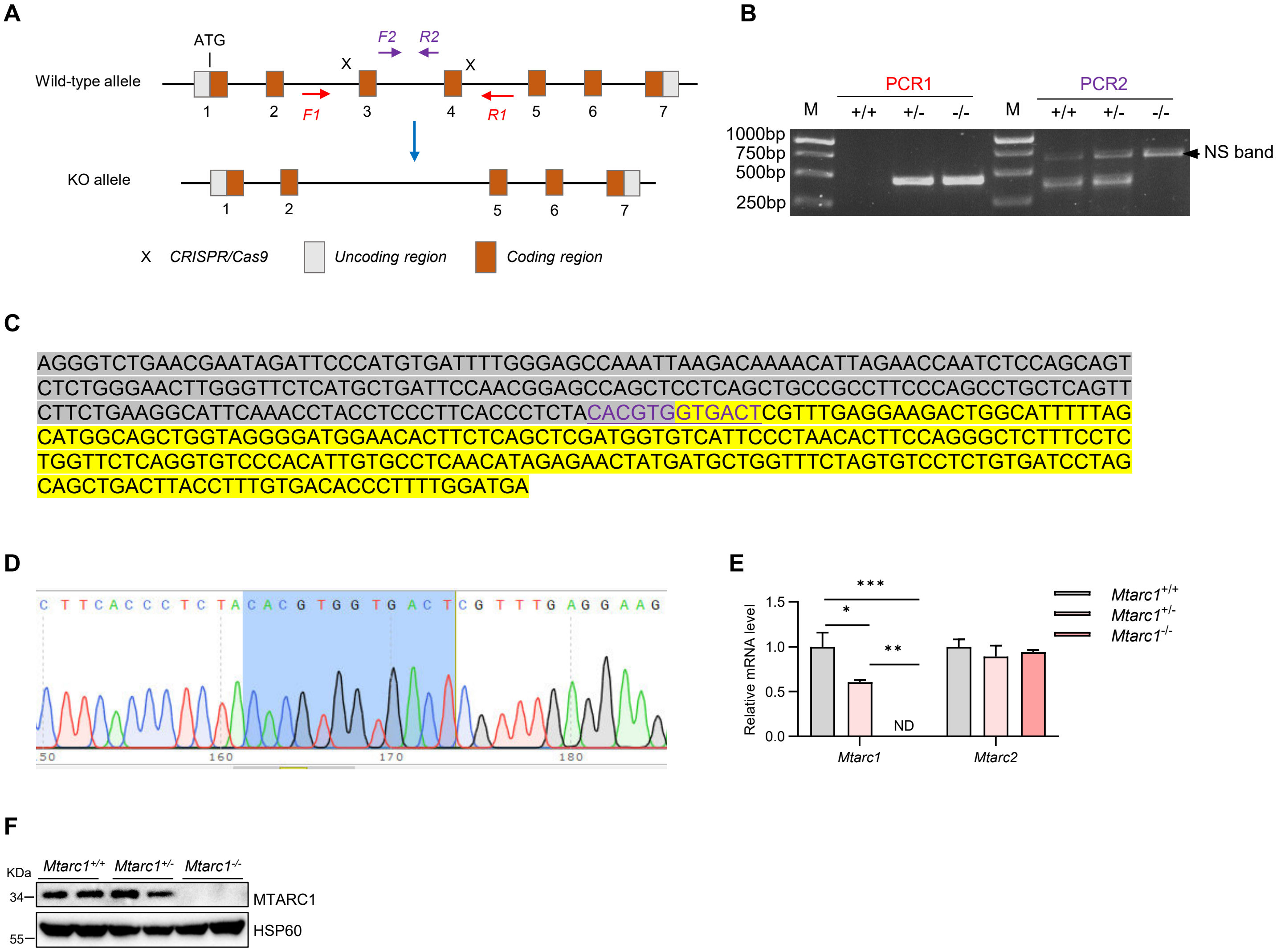


**Fig S1. The generation of *Mtarc1* whole-body knockout mice (related to Fig 1).** (A) The graphic view showing the binding sites of primers for genotyping. F1 and R1 are the primers for PCR1, and the amplicon for knockout and wild-type alleles are 416 bp and 3,520 bp, respectively. The 3,520 bp wild-type band is absent due to the limitation of the extension time of PCR. F2 and R2 are the primers for PCR2, the amplicon for wild type allele is 361 bp, and no amplicon for the knockout allele due to the absence of a template DNA region. (B) Agarose gel showing the genotyping PCR products for *Mtarc1*$^{+/+}$, *Mtarc1*$^{+/-}$, and *Mtarc1*$^{-/-}$. NS band, a non-specific band around 800 bp. (C) The knockout band of PCR1 as shown in Panel B was gel purified and sequenced using primer F1 and R1. The sequences highlighted in grey and yellow are the upstream and downstream flanked sequences of the deleted region, respectively. (D) Twelve nucleotides underlined in Panel C were presented in the sequence file. (E) qPCR detecting the knockout efficiency in livers from *Mtarc1*$^{+/+}$, *Mtarc1*$^{+/-}$ and *Mtarc1*$^{-/-}$. (F) Immunoblotting for MTARC1 KO efficiency. Data were expressed as mean ± SEM and analyzed by Student's t-test. * $p < 0.05$, ** $p < 0.01$, *** $p < 0.001$.

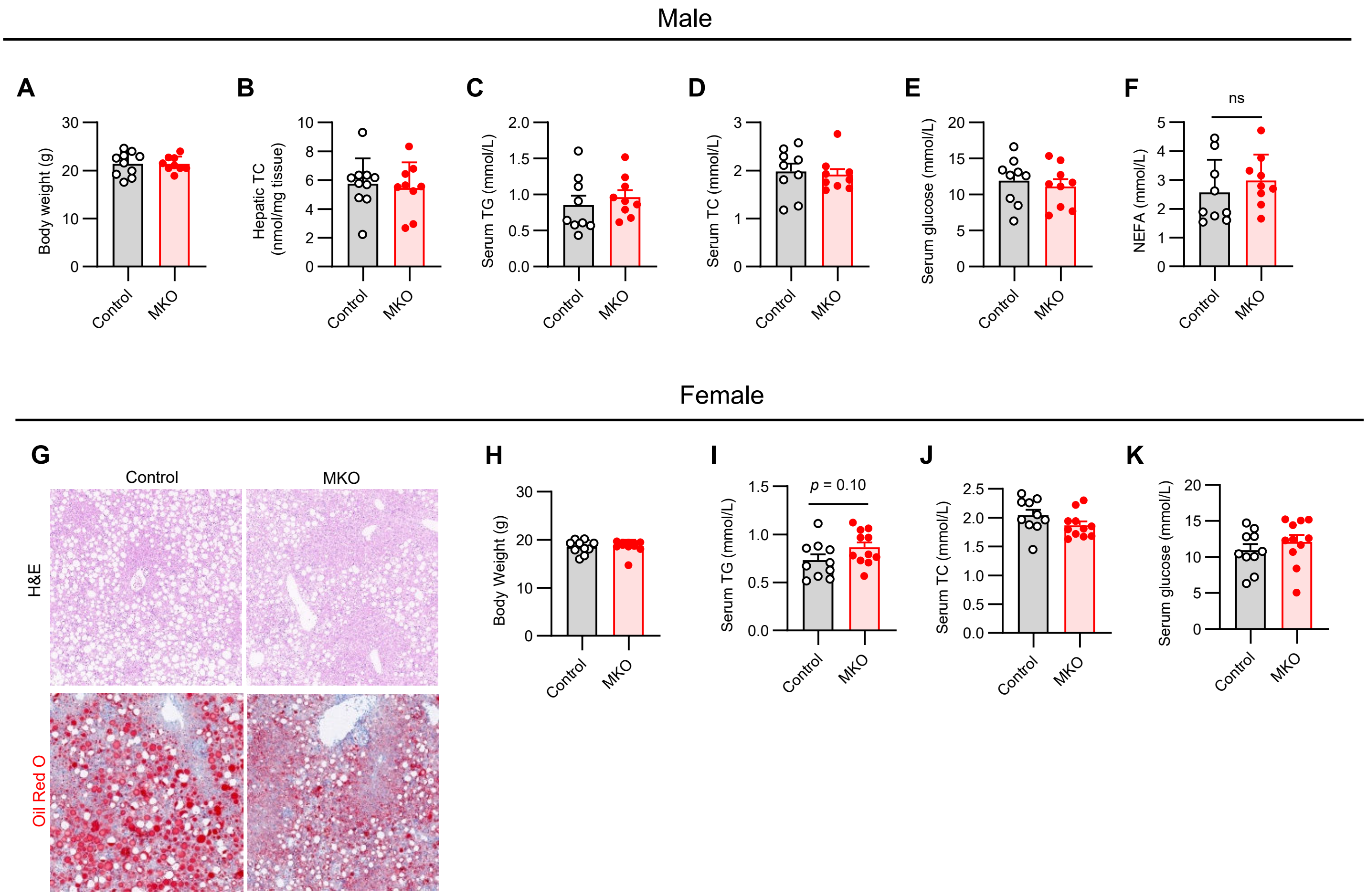


**Fig S2. *Mtarc1* knockout mice were challenged with CDAHFD (related to Fig 1).** (A-F) Male control and *Mtarc1* null mice (MKO) were challenged with CDAHFD for eight weeks and sacrificed for phenotyping (n = 9-10/group). (A) Body weights. (B) Hepatic cholesterol content. (C-D) Plasma triglyceride and total cholesterol content. (E) Serum glucose content. (F) Serum non-esterified fatty acid content. (G-L) Female control and *Mtarc1* null mice (MKO) were challenged with CDAHFD for eight weeks and sacrificed for phenotyping (n = 10/group). (G) Representative images of H&E staining and Oil Red O staining. (H) Body weights. (I-J) Plasma triglyceride and total cholesterol content. (K) Plasma glucose content. Data were expressed as mean ± SEM and analyzed by Student's t-test. * $p < 0.05$, ** $p < 0.01$, *** $p < 0.001$.

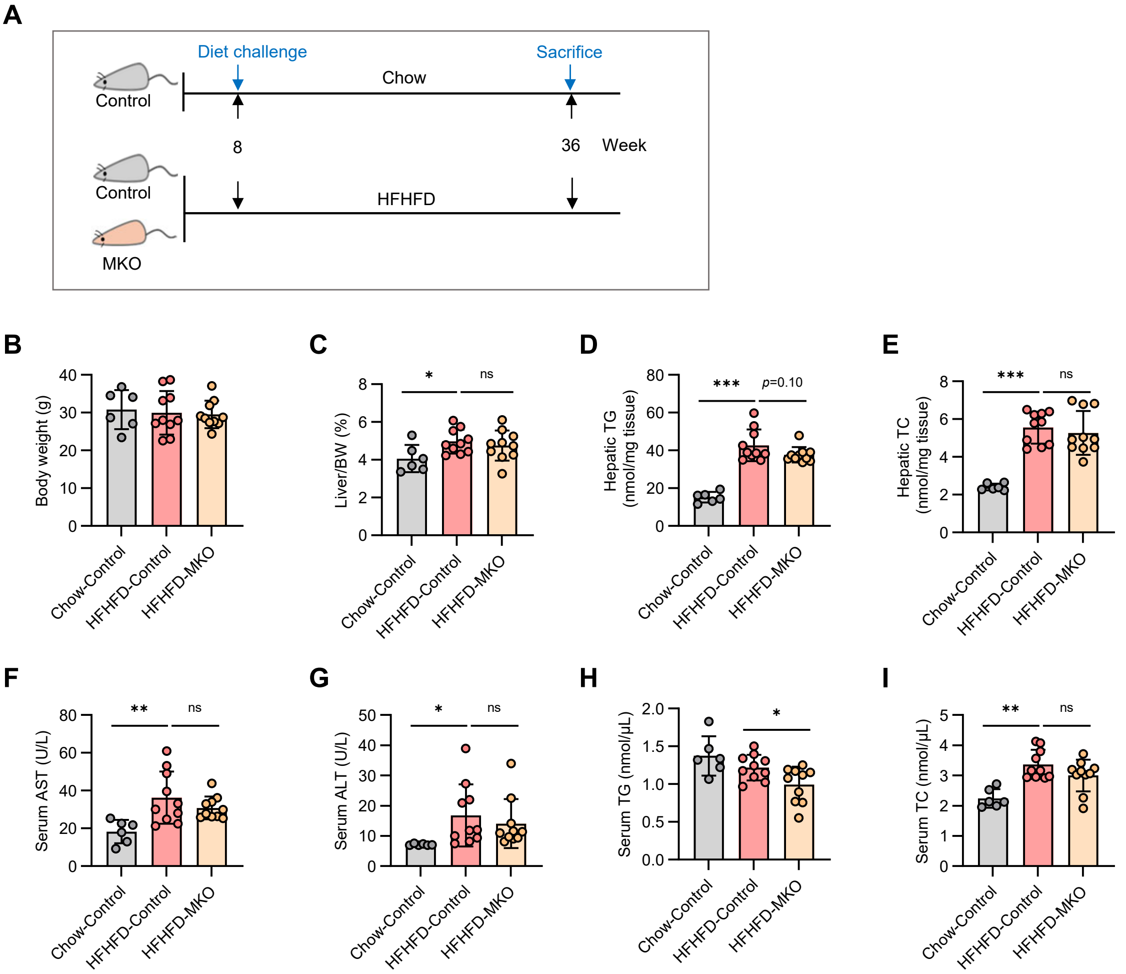


**Fig S3. *Mtarc1* knockout mice were challenged with HFHFD (related to Fig 1).** (A) The graphic view of 8-week-old female control and *Mtarc1* null mice (MKO) were challenged with HFHFD for twenty-eight weeks. One extra control group fed on regular chow was set. (B) Body weights. (C) Liver body weight ratio (%). (D-E) Hepatic triglyceride and total cholesterol content. (F-G) Activity of serum alanine aminotransferase (ALT) and serum aspartate aminotransferase (AST). (H-I) Plasma triglyceride and total cholesterol content. Data were expressed as mean ± SEM and analyzed by Student's t-test. * $p < 0.05$, ** $p < 0.01$, *** $p < 0.001$.

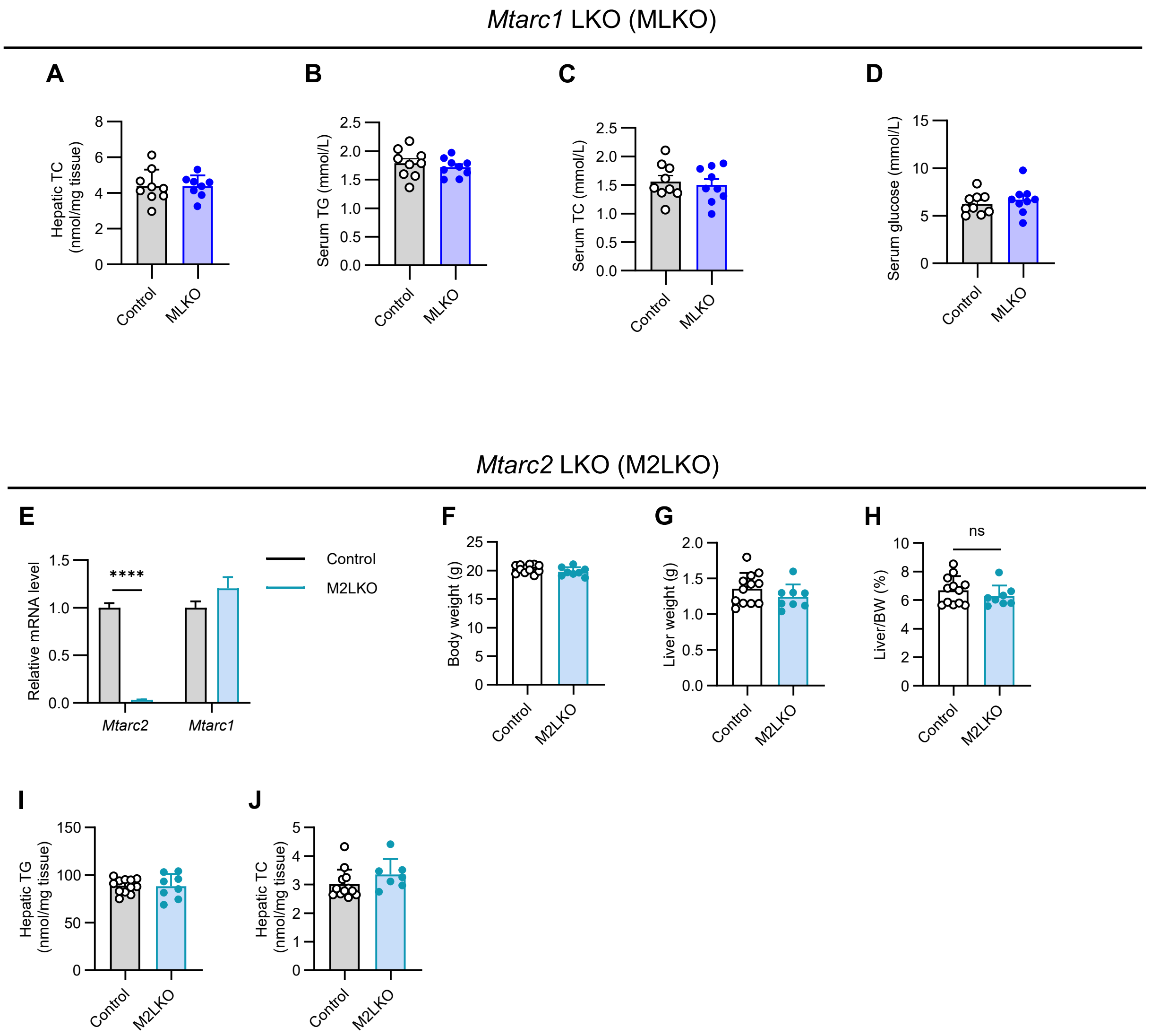


**Fig S4. related to Fig 2.** (A-D) *Mtarc1* liver-specific knockout mice were challenged with CDAHFD. (A) Hepatic cholesterol content. (B-C) Plasma triglyceride and total cholesterol content. (D) Plasma glucose content. (E-J) LSL-Cas9 adult mice were tail vein injected with AAV8 carrying either U6.*Mtarc2*-sg1.U6.*Mtarc2*-sg2.TBG.Cre or non-specific control sgRNA.TBG.Cre. Control and M2LKO mice were challenged with CDAHFD for seven weeks and sacrificed for phenotyping (n = 8-12/group). (E) Hepatic qPCR assay for *Mtarc2* and *Mtarc1*. (F) Body weights. (G) Liver weights. (H) Liver body weight ratio (%). (I-J) Hepatic triglyceride and cholesterol content. Data were expressed as mean ± SEM and analyzed by Student's t-test. **** $p < 0.0001$; ns, no significance.

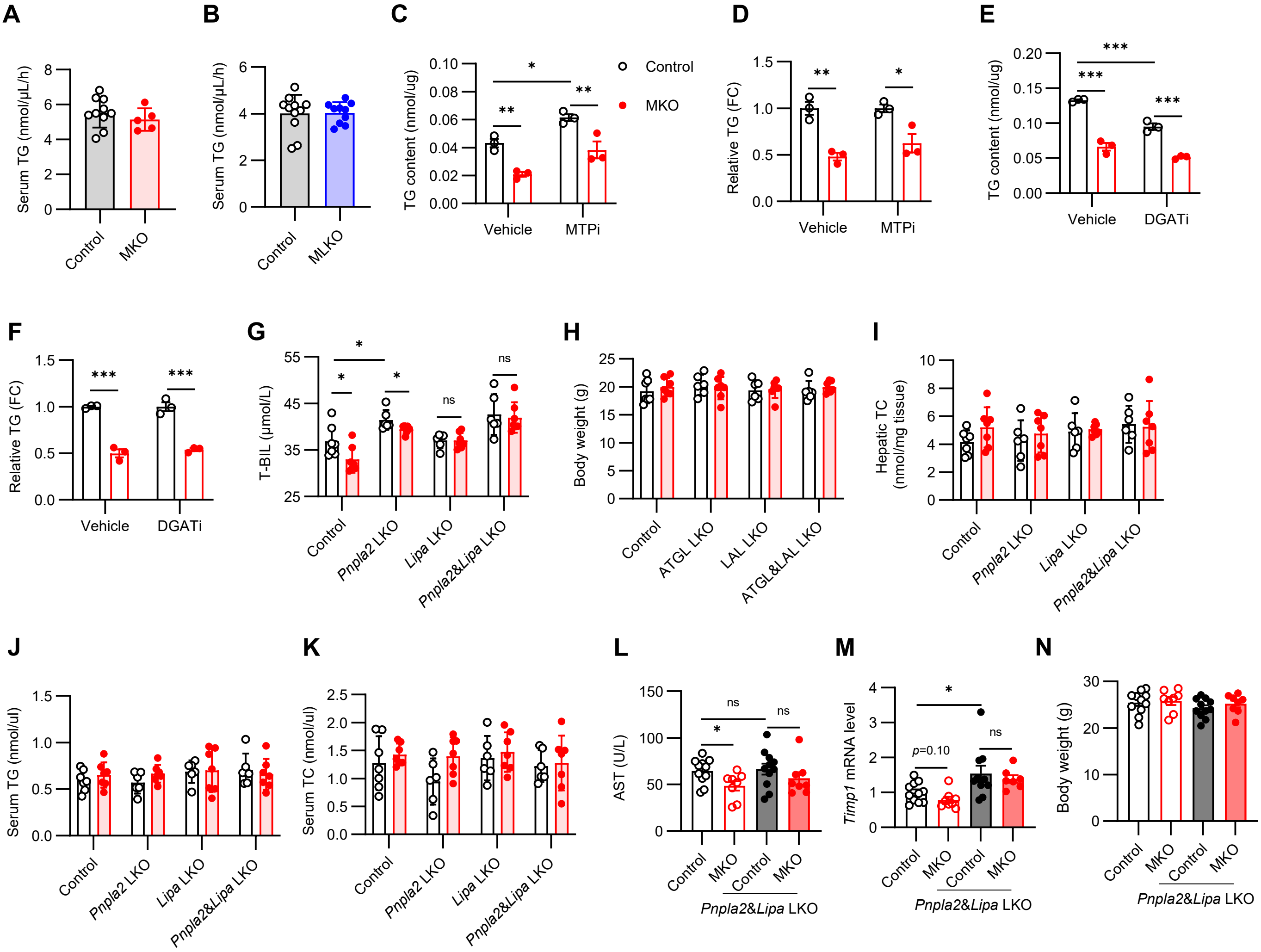


**Fig S5. related to Fig 3.** (A) Male control and MKO mice were fed with CDAHFD for 6 weeks and then subjected to TG secretion assay (n = 5-10/group). (B) Control and MLKO mice were fed with CDAHFD for 5 weeks and then subjected to TG secretion assay (n = 10 to 11/group). (C) Cellular TG content of primary hepatocytes treated with DGATi (TG synthesis inhibitor) as indicated for 24 hours (n = 3/group). (D) The relative TG levels of panel C (n = 3/group). (E) Cellular TG content of primary hepatocytes treated with MTPi (VLDL assembly and transport inhibitor) as indicated for 24 hours (n = 3/group). (F) The relative TG levels of Panel E (n = 3/group). (G-K) related to Fig 3C to I. (G) Serum total bilirubin content. (H) Body weights. (I) Hepatic cholesterol content. (J) Serum triglyceride content and (K) total cholesterol content. (L-N) related to Fig 3J to N. (L) The activity of serum aspartate aminotransferase (AST). (M) The expression level of hepatic *Timp1* assayed by qPCR. (N) Body weights. Data were expressed as mean ± SEM and analyzed by Student's t-test. * $p < 0.05$, *** $p < 0.001$, ns, no significance.

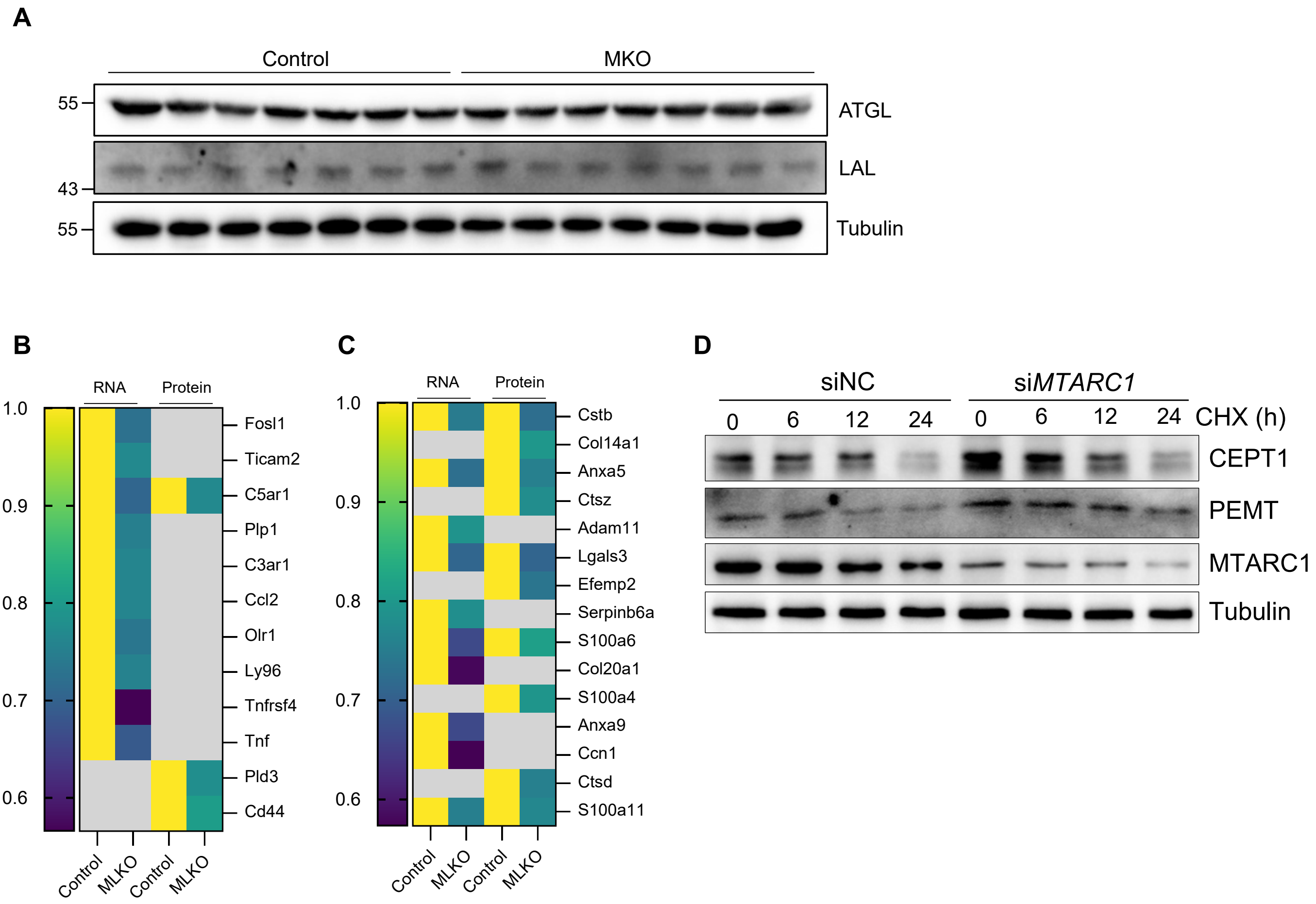


**Fig S6. Hepatic proteomic and transcriptomic data (related to Fig 3)**. (A) Immunoblotting for control and MKO livers (n=7/group). (B) Heatmap presenting the detailed expression profile of the term "inflammatory response" from BP group of the top four terms from gene ontology annotation for downregulated hits. (C) Heatmap presenting the detailed expression profile of the term "collagen-containing extracellular matrix" from CC group of the top four terms from gene ontology annotation for downregulated hits. (D) HepG2 cells were transfected with either nontarget control siRNA or *MTARC1* siRNA. After 36 hours post-transfection, the cells were treated with 300 μM for the duration as indicated.

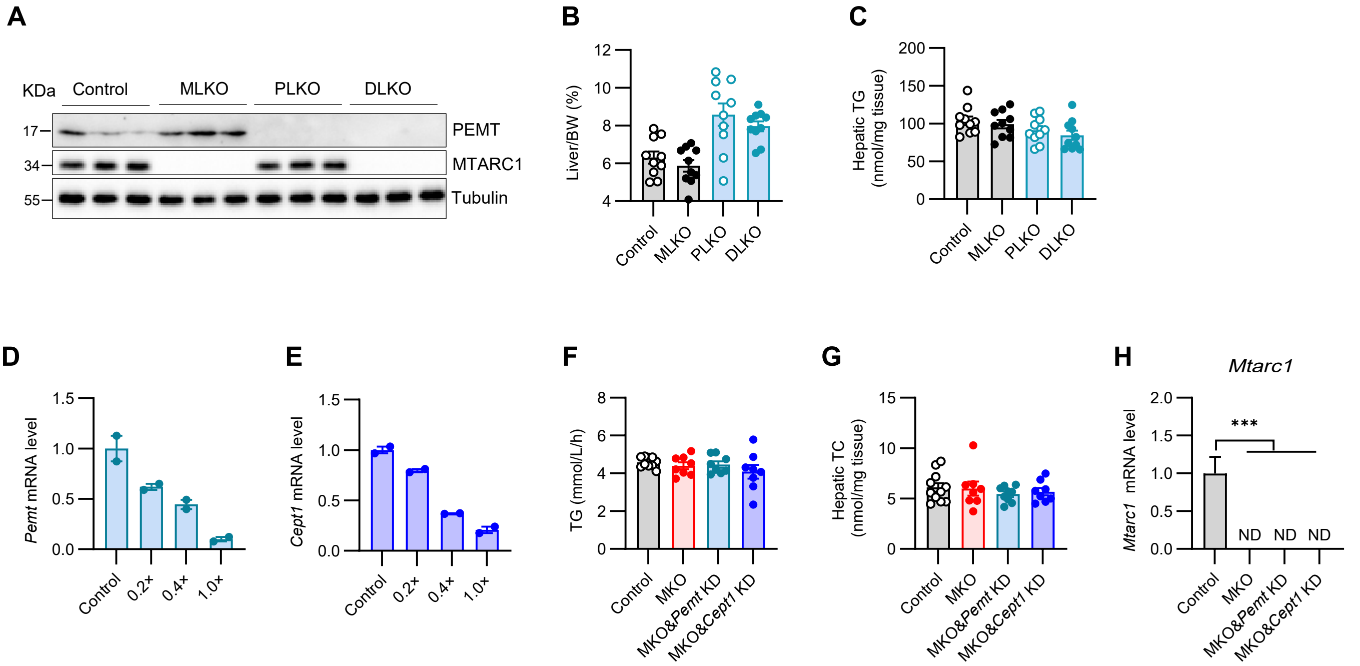


**Fig S7. Related to Fig 6.** (A) Hepatic immunoblotting assay verifying the knock-out efficiency of *Mtarc1* and *Pemt*. (B) Liver body weight ratio (%) (n = 10/group). (C) Hepatic triglyceride content (n = 10/group). (D) Wild-type C57 mice were administrated with AAV8 expressing *Pemt* shRNA at the dosage as indicated (1.0× denotes 1e$^{11}$ vg/mouse) and sacrificed for hepatic qPCR assay after one week (n = 2/group). (E) Wild-type C57 mice were administrated with AAV8 expressing *Cept1* shRNA at the dosage as indicated (1.0X denotes 1e$^{11}$ vg/mouse) and sacrificed for hepatic qPCR assay after one week (n = 2/group). Note, to ensure that an equal amount of virus was used for each mouse, the control virus was used as filler. (F) Plasma VLDL secretion rate. (G) Hepatic cholesterol content. (H) qPCR assay confirming the genotype of *Mtarc1*. Data were expressed as mean ± SEM and analyzed by Student's t-test. *** $p < 0.001$.
